# KDM6A Regulates Immune Response Genes in Multiple Myeloma

**DOI:** 10.1101/2024.02.12.579179

**Authors:** Daphne Dupéré-Richer, Alberto Riva, Sayantan Maji, Benjamin G. Barwick, Heidi Casellas Román, Amin Sobh, Gabrielle Quickstad, Jianping Li, Umasankar De, Crissandra Piper, Marta Kulis, Teresa Ezponda, José Ignacio Martin-Subero, Giovanni Tonon, Weizhou Zhang, Constantine S Mitsiades, Lawrence H Boise, Richard L. Bennett, Jonathan D. Licht

**Author notes:** Corresponding Author: Jonathan D. Licht, MD, The University of Florida Health Cancer Center Cancer/Genetics Complex, 2033 Mowry Road, Suite 145, Gainesville, FL 32610.

## Abstract

The histone H3K27 demethylase KDM6A is a tumor suppressor in multiple cancers, including multiple myeloma (MM). We created isogenic MM cells disrupted for KDM6A and tagged the endogenous protein to facilitate genome wide studies. KDM6A binds genes associated with immune recognition and cytokine signaling. Most importantly, KDM6A binds and activates *NLRC5* and *CIITA* encoding regulators of Major Histocompatibility Complex (MHC) genes. Patient data indicate that NLRC5 and CIITA, are downregulated in MM with low KDM6A expression. Chromatin analysis shows that KDM6A binds poised and active enhancers and KDM6A loss led to decreased H3K27ac at enhancers, increased H3K27me3 levels in body of genes bound by KDM6A and decreased gene expression. Reestablishing histone acetylation with an HDAC3 inhibitor leads to upregulation of MHC expression, offering a strategy to restore immunogenicity of KDM6A deficient tumors. Loss of *Kdm6a* in murine RAS-transformed fibroblasts led to increased growth *in vivo* associated with decreased T cell infiltration.

**Statement of significance:** We show that KDM6A participates in immune recognition of myeloma tumor cells by directly regulating the expression of the master regulators of MHC-I and II, NLRC5 and CIITA. The expression of these regulators can by rescued by the HDAC3 inhibitors in KDM6A-null cell lines.

## Introduction

Multiple myeloma (MM), the second most common hematological cancer, is a malignant proliferation of immunoglobulin-secreting plasma cells. Treatment of this disease has undergone significant advances with the addition of immunomodulators, proteasome inhibitors, anti-tumor antibodies and most recently engineered T cell therapy. Despite this progress, MM remains an incurable disease and accounts for 20% of all deaths from hematological cancers. The molecular genetics of MM was initially elucidated by profiling cytogenetic (1–3), gene expression (4,5) and gene copy number changes in tumor specimens (6). These studies indicated that deregulation of transcription factors (MAF, IRF4, MYC) and cofactors (NSD2) is common in MM, and often due to rearrangement with the immunoglobulin enhancers. More recently, next generation sequencing surveys indicated the presence of mutations affecting a range of epigenetic regulator genes in up to 25% of MM patients [4]. Among the affected genes are those encoding the histone demethylase (HDM) KDM6A, the histone methyltransferases (HMT) KMT2C and KMT2D, the histone acetyltransferase (HAT) CREBBP and the chromatin remodeling subunit ARID1A. These proteins interact together to regulate enhancers (7), emphasizing the importance of enhancer deregulation in MM.

*KDM6A* undergoes inactivating mutations and deletions across many tumor types with an incidence of 4% in the cBioPortal (8). Across all tumor types KDM6A mutation is associated with a >40% decrease in median survival time (8). *KDM6A* is present on the X chromosome and is expressed from both X alleles, while in men the homologous UTY gene is expressed alongside with UTX from a single gene. Of note, the X chromosome is commonly lost in MM cases, according to cBioPortal data occurring in MM reaching 16% females (8). *KDM6A* mutation or deletion occurs in ∼3% of MM cases at diagnosis and was associated with poor survival (9). Notably, more than one third of all MM cell lines have *KDM6A* anomalies, and these cells were generally derived from advanced cases of MM, suggesting that KDM6A loss may be a progression factor in MM. In agreement with this idea, we showed that KDM6A acts as a tumor suppressor in MM by modulating cell growth and adhesion (10).

The major KDM6A isoform is 1400 amino acids long and contains tetratricoid repeats involved in protein-protein interactions (11) and a C-terminal JmJC demethylase domain that removes methyl groups from histone H3 at lysine 27 (H3K27) in a Fe^2+^ –dependent reaction, requiring α-ketoglutarate as a cofactor. KDM6A interacts and works in concert with the COMPASS-like complex which includes the H3K4me1 methyltransferases KMT2C or KMT2D. KDM6A helps activate enhancers by removing the repressive H3K27me3 chromatin modification and recruiting P300 or CREBBP histone acetyltransferases to acetylate H3K27 at active enhancers (12). The EZH2 methyltransferase component of the PRC2 complex, that catalyzes the H3K27 trimethylation reaction, is often overexpressed in MM, and also correlates with poor prognosis (17). Chromatin profiling studies in primary MM (15) also support the concept that an imbalance in epigenetic regulation at enhancers may contribute to the biology of MM. We found that PRC2/EZH2 inhibition can compensate for the loss of KDM6A, suggesting that the catalytic activity of KDM6A plays a role in tumor suppression. On the other hand, KDM6A tumor suppressive functions have been linked to its ability to interact with transcription factors and other epigenetic regulators. Indeed, mutation in the TPR domains or deletion of the TPR of KDM6A compromises interaction with the KMT2C/D COMPASS-like complex (16) and abolishes growth suppression by KDM6A (17). Furthermore, the recruitment of histone acetyltransferases can be achieved by KDM6A mutant devoid of HDM activity (18).

Aiming to understand the function of KDM6A in MM, we used CRISPR engineered isogenic cell lines and ChIP-sequencing to identify KDM6A binding sites in the myeloma genome. We further explored how KDM6A deficiency affects chromatin structure at these loci by mapping genome-wide histone modifications. We found KDM6A bound at enhancers in proximity to genes regulating immune function, notably the master regulators of class I and II MHC, NLRC5 and CIITA. Furthermore, depletion of KDM6A leads to a decrease in H3K27ac at KDM6A associated enhancers while H3K27me3 is decreased in the body of genes where the promoter or the enhancer is bound by KDM6A. We found a correlation between expression of KDM6A and CIITA and NLRC5 in MM patients. Modeling KDM6A loss *in vivo* revealed impaired T cell infiltration in tumors where *Kdm6a* is depleted, suggesting that KDM6A regulates tumor immunogenicity.

## Results

### Loss of KDM6A confers a transient growth advantage to multiple myeloma cells

We previously demonstrated the tumor suppressive function of KDM6A in MM cell lines in vitro and in vivo using an add-back system (10). To further corroborate our previous findings, we examined expression of KDM6A in 764 newly-diagnosed MM patients from the CoMMpass trial. Consistent with tumor suppressive functions of KDM6A, we found longer progression free survival (PFS) in female patients within the high KDM6A expression quartile (**Fig. 1A**). However, in male patients, which express lower levels of KDM6A compare to female patients (**Fig. 1B**), there is no correlation between KDM6A expression and PFS. This suggested that KDM6A is a sex-specific tumor suppressor, therefore we continued experiments in female cell lines.

**Figure 1:**
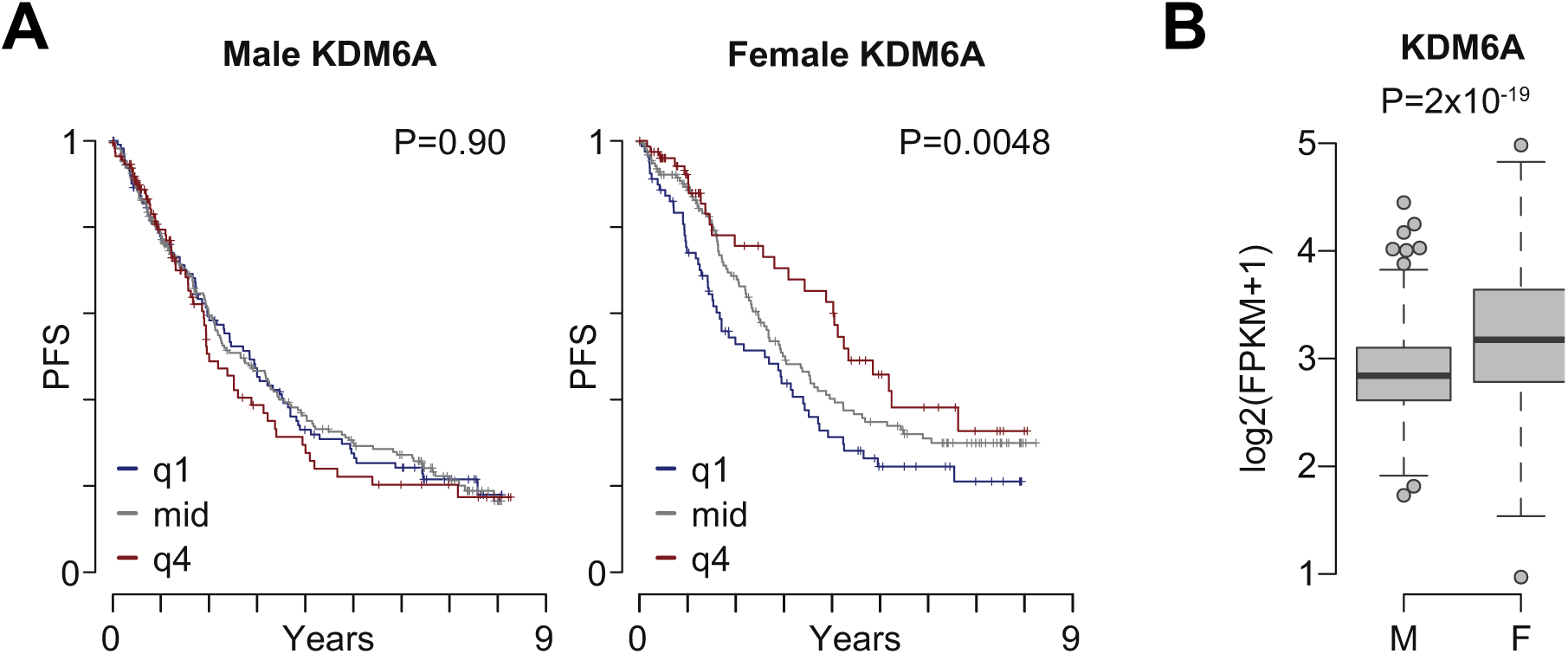
KDM6A has properties of a tumor suppressor in female myeloma patients. **A**) Kaplan-Meier curves of male (left) and female (right) CoMMpass patients stratified by KDM6A expression into low (q1; blue), two mid (gray), and high (q4; red) quartiles. P-values indicate the cox-proportional hazards regression P-value of KDM6A expression associated with progression-free survival (PFS). **B)** Expression of KDM6A in 764 newly-diagnosed MM patients from the CoMMpass trial. P-value determined by Wilcoxon test.

To confirm the consequences of KDM6A loss in MM, we transiently transfected female myeloma cell lines Karpas-620, AMO1 and MM.1S with a ribonucleoprotein complex of CAS9 and KDM6A specific guide RNAs. Gene editing of exon 4 was efficient with 40-60% of cells displaying predominantly 1, 2 and 5 bp deletions and 1bp insertions (supplemental table S1) within 2 days of transfection. By 8 days of growth this number increased to 80% implying a growth advantage to cells devoid of KDM6A (supplementary Fig.S1A). Gene editing with guides targeting exons 1, 4 or 6 yielded cells pools with dramatically reduced or absent levels KDM6A protein in the absence of selection suggesting a growth advantage to cells that acutely lose KDM6A expression (**Supplementary Fig. S1B**). However, after a few weeks in culture, KDM6A depleted cell pools did not exhibit a proliferation advantage over a WT cell pool that was electroporated with a non targeting gRNA (Supplemental Fig. S1C). This suggests that KDM6A may have growth suppressive functions but cells can adapt and reach a new homeostatic state.

### KDM6A binds genes that regulates the immune system

To determine direct targets of KDM6A in MM we used ChIPmentation, which combines chromatin immunoprecipitation and cleavage of chromatin with a transposase (19), to identify KDM6A binding sites in two female myeloma cell lines, ARP1 and Karpas-620 which express wild type KDM6A (Supplemental Fig. S1B and (20)).In ARP1 cells we identified 28,119 peaks that were absent in KDM6A-null ARP1 E4 cells. To validate KDM6A binding sites and the quality of the KDM6A antibody, we inserted a dual HA tag sequence into the C-terminal of KDM6A by CRISPR-mediated gene editing of ARP1 cells (**Fig. 2A**). Using the HA antibody, we detected 13,251 binding sites overlapping with the those detected with an antibody against KDM6A (**Fig. 2B and supplemental Table S2)**. These binding sites are mainly found in intergenic regions (62%) and introns (24%) (**Fig. 2C**), consistent with the localization of KDM6A to enhancers. With the use of Genomic Regions Enrichment of Annotations Tool (GREAT version 4.0.4.) (21) we linked these peaks to 3863 genes at distances <100kb. However, most of the peaks (8371) were further than 100kb from any gene (**Fig. 2D**). To obtain a functional understanding of KDM6A regulated network in myeloma, we performed pathway enrichement analysis of the genes found less than 100kb away from KDM6A binding sites using ShinyGo v0.77 (22), using a background reference of all genes expressed in the ARP1 cells (FPKM >10).

**Figure 2:**
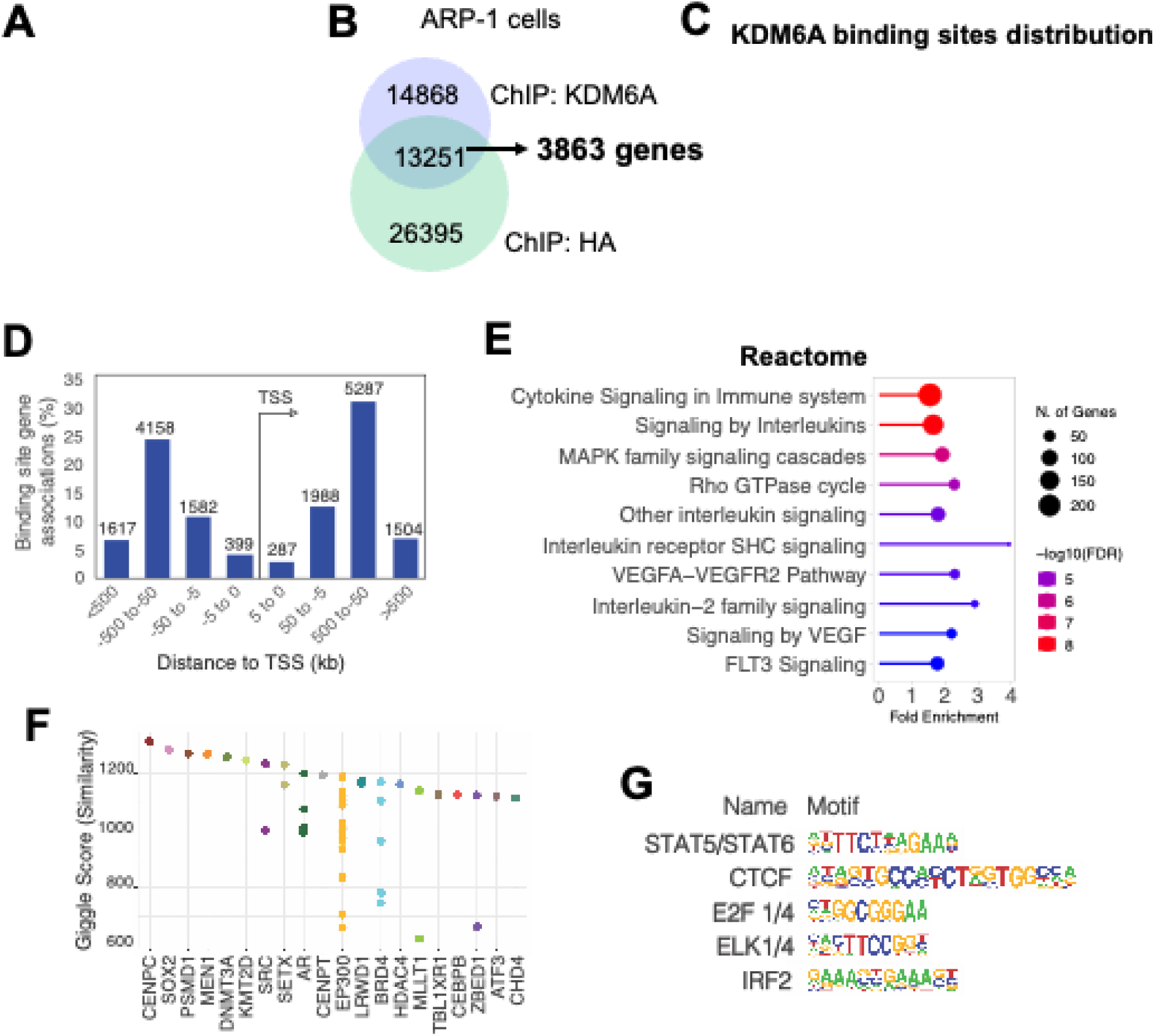
KDM6A binding sites in myeloma cell lines. **A**) Immunoblot showing the detection of KDM6A by an HA antibody in ARP1 cells in which both alleles of endogenous KDM6A was HA-tagged using CRISPR-Cas9 gene editing. ARD is a KDM6A negative control cell line. **B)** Overlap of KDM6A binding sites detected by chromatin precipitation of an HA tagged ARP1 cell line with anti-HA or anti-KDM6A antibodies. **C)** Distribution of KDM6A binding sites in the annotated regions of the genome in the ARP1 cell line. **D)** Distance and orientation between KDM6A binding regions and their closest genes. **E)** Dot plot visualization of GO biological process enrichment analyses of KDM6A bound genes using the GoShiny v0.77 tool. The size of the dots reflects the number of genes, the length of the line with fold enrichment and color scale indicates false discovery rate. **F)** Similarity of KDM6A binding pattern with those of transcription factors found in the Cistrome data base (top 20). Each dot represents a different ChIP experiment. **G)** HOMER-identified enriched transcription factor binding motif within regions bound by KDM6A as identified both by KDM6A and HA antibody in ARP1 cells expressing HA-tagged KDM6A.

Consistent with our previous findings, we observed that many bound genes are associated with cell signaling, motility and adhesion. Interestingly however, most of the KDM6A bound genes are related to the regulation of immune system (**Fig. 2E)**. Among the genes in close proximity to KDM6A binding sites we found the transcriptional regulators of class I Major histocompatibility complex (MHC) genes, NOD-like receptor family CARD domain containing 5 (NLRC5), and of class II (MHC-II), MHC class II transcriptional activator (CIITA). Comparing the KDM6A binding pattern to previously reported ChIP-Seq experiments reported in the cistrome database (23) we found similarity of KDM6A binding to that of KMT2D and EP300, supporting a potential functional interaction of KDM6A with the COMPASS-like complex in myeloma cells (**Fig. 2F)**. Motif enrichment analysis of the genomic region bound to KDM6A using the HOMER package (24), revealed significant enrichment of binding sites for CTCF, E2F1/4 and the ETS related ELK1/4 as well as IRF2 and STAT5 both effectors of immune signaling (**Fig. 2G and Supplemental Table S3**). To corroborate these findings, ChIP-Seq analysis of Karpas-620 cells with KDM6A antibody revealed over 60,000 peaks, 6163 overlapping with the highly validated peaks found by both HA and KDM6A antibodies in ARP-1 cells (**Supplementary** Fig. 2A). These binding sites were found to be within 100kb of 2,268 genes involved in the immune system, response to stimulus and cell motility (**Supplementary** Fig. 2B). Motif enrichment analysis identified the same transcription factor recognition sites as those found in the ARP1 cell line i.e., CTCF, STAT5, E2F1 and IRF2 (**Supplementary** Fig. 2C **and Supplemental Table S3**).

### Loss of KDM6A is associated with decreased expression of genes regulating immunity

By CRISPR mediated gene editing we generated isogenic clonal cell lines with and without *KDM6A* and analyzed the downstream impact on the transcriptome. In the clonal EJM cell lines we used a guide RNA directed to exon 6 of KDM6A to generate a homozygous knockout. In the AMO-1 cells we created a heterozygous KDM6A mutant clonal cell lines using a gRNA against exon 1 (**Fig. 3A**). There was no consistent pattern of gene upregulated with KDM6A loss across the cell lines (**Supplementary Fig. S3A right panel**). There was a more consistent overlap of genes downregulated by KDM6A KO in the 3 cell lines including MHC-II genes. More specifically we found HLA-DRB1, HLA-DPA1, HLA-DR, HLA-DPB1 and the master regulator of MHC-II gene expression, CIITA (**Supplementary Fig. S3A and Supplemental Table S4**). A heatmap representation of RNA-Seq data (z-score) showed decreased expression of most MHC-II genes in AMO and EJM clonal cell lines depleted for KDM6A, and MHC class I genes were downregulated in KDM6A null ARP1 cells (**Fig. 3B**). Flow cytometry analysis confirmed significant downregulation of MHC-II, HLA-DR/DQ/DP expression at the cell surface of AMO and EJM KDM6A KO cells (**Fig. 3C**) while ARP1 cells do not express detectable MHC II. Similarly, MHC I (HLA-A/B/C) expression decreased when KDM6A was eliminated in ARP1 and AMO-1 cell lines. ChIP experiments revealed that KDM6A bound to NLRC5 and CIITA (**Fig. 3D**), and we confirmed that both genes were significantly downregulated at the mRNA level in KDM6A-depleted clonal cell lines (**Fig. 3E**), suggesting a direct role for KDM6A in regulation of gene transcription. We reanalyzed the data of the Multiple Myeloma Research Foundation CoMMPass comprising 694 male and 460 female patients in which KDM6A expression levels were available. Using the top and bottom 10^th^ percentile as a cut-off for high and low expression of KDM6A, respectively we observed that NLRC5 and CIITA, were significantly decreased in both male and female KDM6A low-expressing tumors (**Fig. 3F**). Furthermore, genes included in the Antigen Presentation Pathway of the gene ontology were downregulated in the low KDM6A group in both male and female patients (**Supplementary** Fig. 3B). Gene expression profiles of MM cells included in the DepMap database (25) show that those in the bottom 10 percentile for KDM6A express lower NLRC5 and CIITA (**Supplementary. Fig. 3C**). Multiplex ELISA assays showed that several cytokines were express at lower levels in the supernatant of KDM6A depleted cells, including the chemotactic cytokines for T-cells, C-C Motif Chemokine Ligand 5 (CCL5) and 7 (CCL7), the inflammatory C-C Motif Chemokine Ligand 3 (CCL3), interferon alpha2 (IFNα2) and the growth factor Fms Related Receptor Tyrosine Kinase 3 Ligand (FLT3L) (**Supplementary** Fig. 3D). Conversely, the mitogenic factor CX3C motif chemokine receptor 1 (CX3CR1) and the platelet derived growth factor subunit A (PDGFA) were elevated in the media of KDM6A depleted cells. Furthermore, transcriptional changes induced by IFNα stimulation in ARP1 cells were attenuated when KDM6A was depleted (**Supplementary** Fig. 4A). Notably, IFNα upregulated MHC-I and II RNA (Supplementary Fig. 4B) and MHC-II protein at the surface of ARP1 cells and this induction is blunted in KDM6A depleted cells (**Supplementary** Fig. 4C).

**Figure 3:**
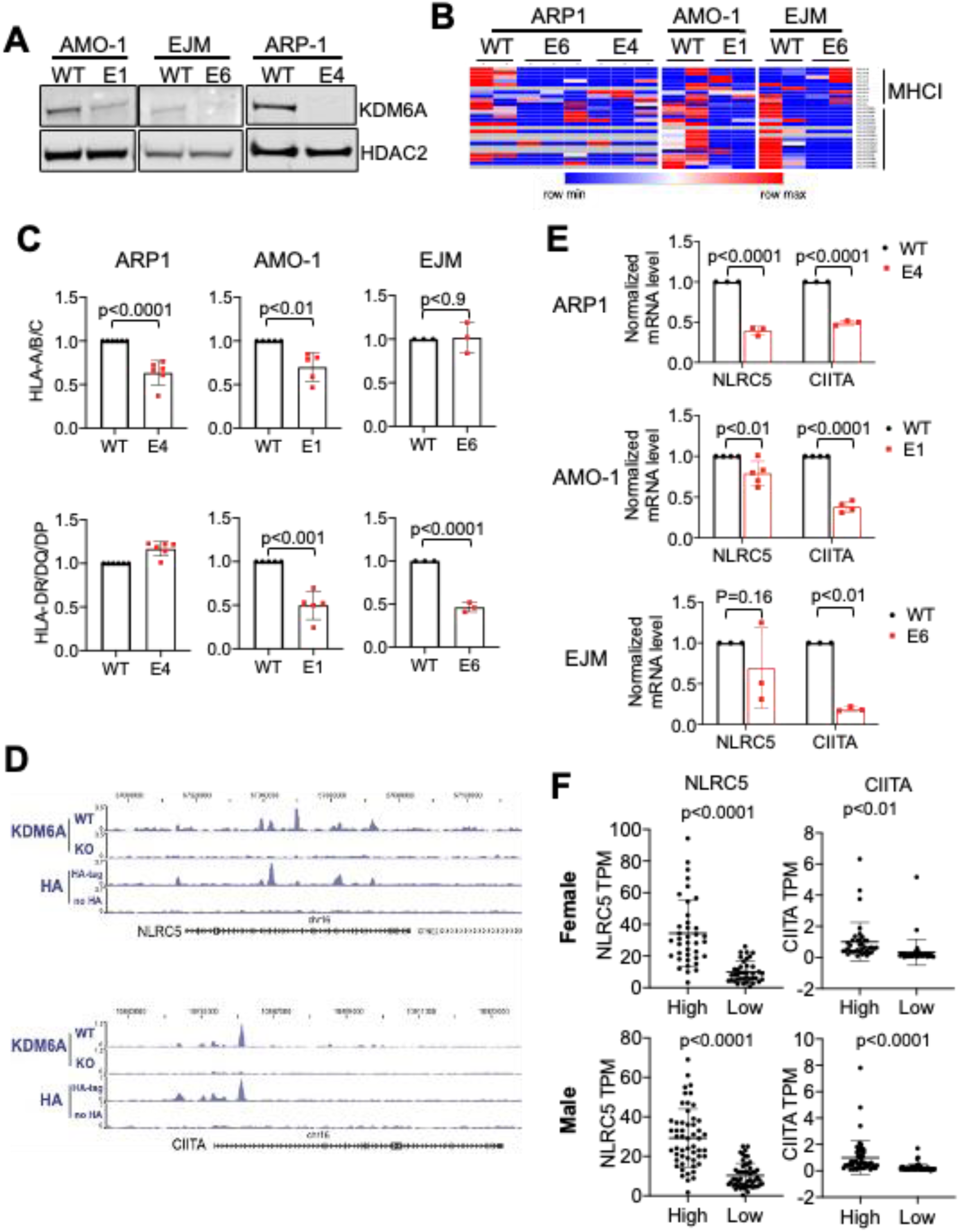
KDM6A controls MHC-I and II gene expression. **A**) Immunoblot showing depletion of KDM6A protein in CRISPR edited clonal cell lines. **B)** Heatmap of z-scores for all expressed MHC genes in KDM6A wild type and knockout cell lines. **C)** Flow cytometry analysis of HLA-A/B/C and HLA-DM/DQ/DR in clonal isogenic cell lines ARP1, AMO-1 and EJM. Bottom panels of each histograms represent the mean fluorescence intensity (MFI) quantification (3 to 6 biological replicates; +/− SD. Wilcoxon T-test). **D)** Genome browser view of the CIITA and NLRC5 locus in ARP1 cell line WT or KO for KDM6A. **E)** mRNA analysis by Q-PCR normalized to GAPDH (3 to 5 biological replicates; +/− SD. Wilcoxon T-test) **F)** Normalized expression values (TPM) of NLRC5 and CIITA in myeloma patient tumors within the top or bottom decile of KDM6A expression (38 females and 54 males in each group, unpaired T-test).

Re-expression of KDM6A was able to induce cell surface expression of MHC II in some KDM6A null cell lines but did not affect MHC I expression (**Supplementary** Fig. 5). The inconsistent rescue of MHC expression with re-expression of KDM6A suggests that a lasting reprogramming of the epigenome occurs in cells after KDM6A loss.

### KDM6A is found at enhancers and regulates H3K27me3 in gene bodies

To understand how KDM6A regulates chromatin to induce transcriptional changes in MM we mapped H3K27ac, H3K27me3 and H3K4me1 in ARP1 clonal cell lines using ChIPmentation and assessed DNA accessibility by ATAC-seq. We defined enhancers as all H3K4me1 peaks greater than 2kb from transcriptional start site while active enhancer had both H3K27ac and H3K4me1. Poised enhancers harbored H3K27me3 and H3K4me1 and inactive enhancers, were marked only by H3K4me1 without H3K27ac or H3K27me3. KDM6A was found associated mainly with inactive enhancers (50% of binding sites) followed by poised enhancers (36%), active enhancers (11%) and very few promoters (2%) (**Fig. 4A**). Genes associated with active enhancers were downregulated in KDM6A deleted cells, reflecting an activating function of KDM6A. Genes associated with inactive and poised enhancers did not show consistent change upon deletion of KDM6A as these genes are generally not expressed (**Supplementary** Fig. 6A). Active gene networks affected by KDM6A loss were involved in lymphocyte activation, immune system development and cell motility (**Fig. 4B**).

**Figure 4:**
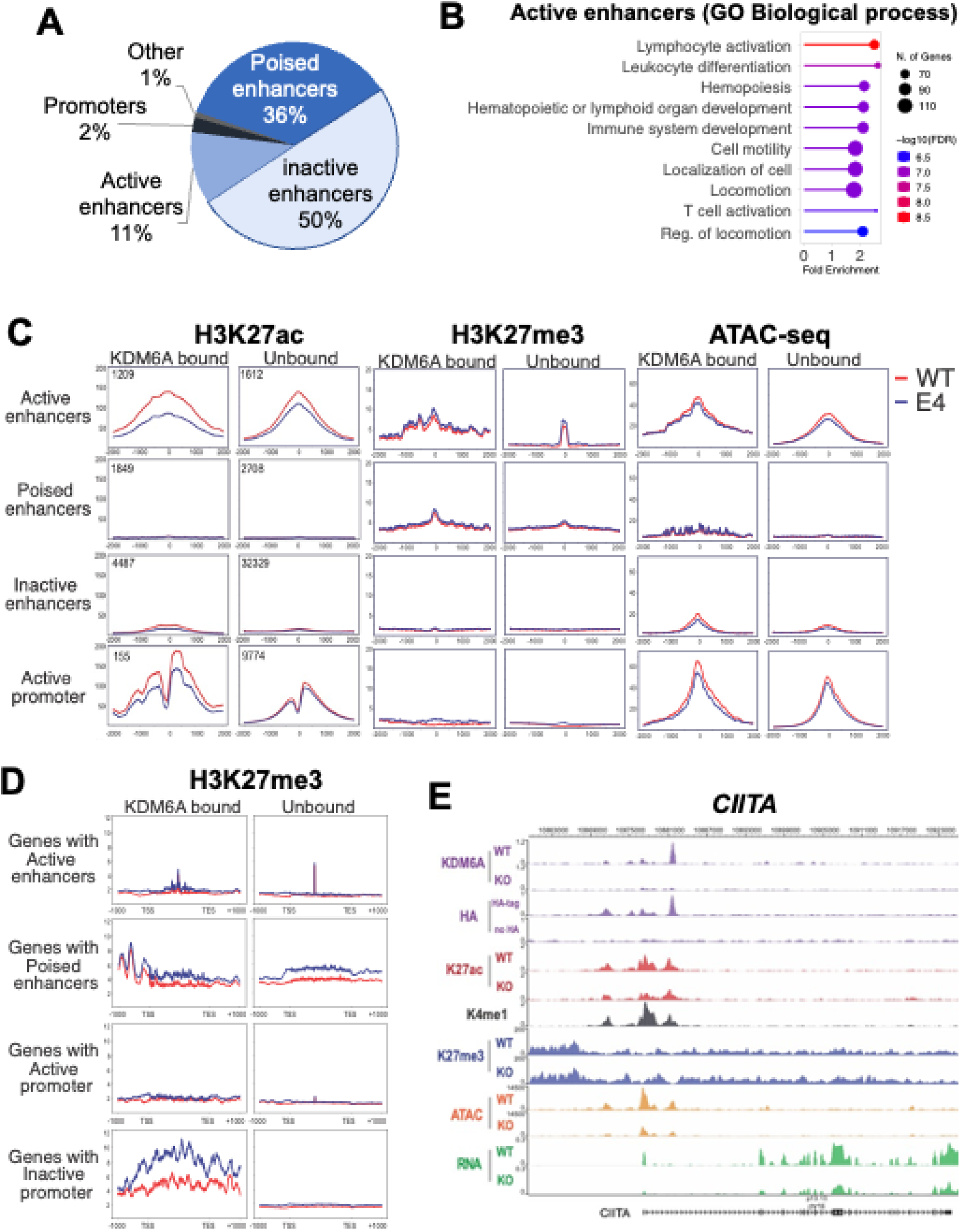
KDM6A loss impact on chromatin structure. **A**) Distribution of KDM6A binding sites annotated regions of the genome in ARP1 cell lines. **B)** Dot plot visualization of GO biological process enrichment analyses genes within 100kb KDM6A-bound enhancers using the GoShiny v0.77 tool. **C)** H3K27ac-seq, H3k27me3-seq and ATAC-Seq signals in ARP1 isogenic clonal cell line WT or KO for KDM6A centered on H3K4me1 peaks for enhancers and centered on transcription start sites for promoters **D)** H3K27me3 metagene analysis at locus bound by KDM6A in ARP1 isogenic clonal cell line WT or KO for KDM6A. **E)** Genome browser view of the CIITA locus in ARP1 cell line WT or KO for KDM6A.

KDM6A KO cells displayed a genome wide decrease in DNA accessibility independent of KDM6A localization (**Supplementary** Fig.6B). KDM6A deficiency caused a reduction in H3K27ac at active enhancers and promoters that it bound. In addition, KDM6A loss led to increased H3K27me3 in wide genomic areas covering the body of genes and found in close proximity to enhancers and promoters bound by KDM6A (**Fig. 4C, D)**. All these changes were represented at both NLRC5 and CIITA loci, where KDM6A was bound at multiple sites in intronic enhancers and promoters (**Fig. 4E** and Supplementary Fig. 6C). Loss of KDM6A at these loci was associated with decreased H3K27ac, chromatin accessibility, and gene expression while H3K27me3 was increased in the vicinity of promoters and within gene bodies.

Using a CRISPR knock in strategy, we created an ARP1 cell line harboring a jmjD-dead KDM6A where 2 amino acids essential for KDM6A enzymatic activity, H1146 and E1148 (26,27), were replaced with alanine residues. ChIP-sequencing indicated that the pattern of H3K27ac modification in jmjD-dead KDM6A expressing cell lines was similar to that of KDM6A wild type cells with a very modest decreased of H3K27ac at active enhancers (Supplementary Fig. 7A). By contrast, there was in an increase in H3K27me3 in the body of genes with inactive promoters in jmjD-dead KDM6A expressing cell similar to that of KDM6A null cells (Supplementary Fig. 7B). This suggest that the tumor suppressive function of KDM6A relies more on its ability to cooperate with other factors to activate genes in association with changes in H3K27Ac rather than its ability to demethylate specific regions.

Studies has shown that KDM6A deficiency can cause a redistribution of its interacting factors leading to enhancer reprogramming in cancer cells and thus activation of a new transcriptional program (28,29). However, ranked ordering of super-enhancers (ROSE) (30) in parental and knockout cells did not indicate enhancer reprogramming upon KDM6A depletion consistent with mild transcriptional changes occurring between KDM6A wild type and depleted ARP1 cells (data not shown). Instead, we observed a global decrease in acetylation at every super enhancer in ARP1 cell when KDM6A is deleted (Supplementary Fig. S7C).

### HDAC3 inhibition restores MHC expression in KDM6A depleted cells

Decreased H3K27ac in KDM6A-null cells suggests that lower activity of CREBBP/EP300 at KDM6A binding sites may lead to decreased expression of KDM6A-associated genes. We previously found that HDAC3 was the enzyme responsible for counterbalancing CREBBP/P300 mediated acetylation of histones in myeloma cell lines (31). Accordingly, treatment of KDM6A knockout ARP1 and AMO-1 MM cells with the HDAC3-selective inhibitor RGFP966 increased total level of acetylated H3K27 (**Fig. 5A)** and upregulated the mRNA for NLRC5 and CIITA (**Fig. 5B**), and their downstream targets, genes encoding MHC-I and II molecules (**Fig. 5C**). Furthermore, RGFP966 restored MHC-I and MHC-II expression at the cell surface of *KDM6A* knockout cells (**Fig. 5D**).

**Figure 5:**
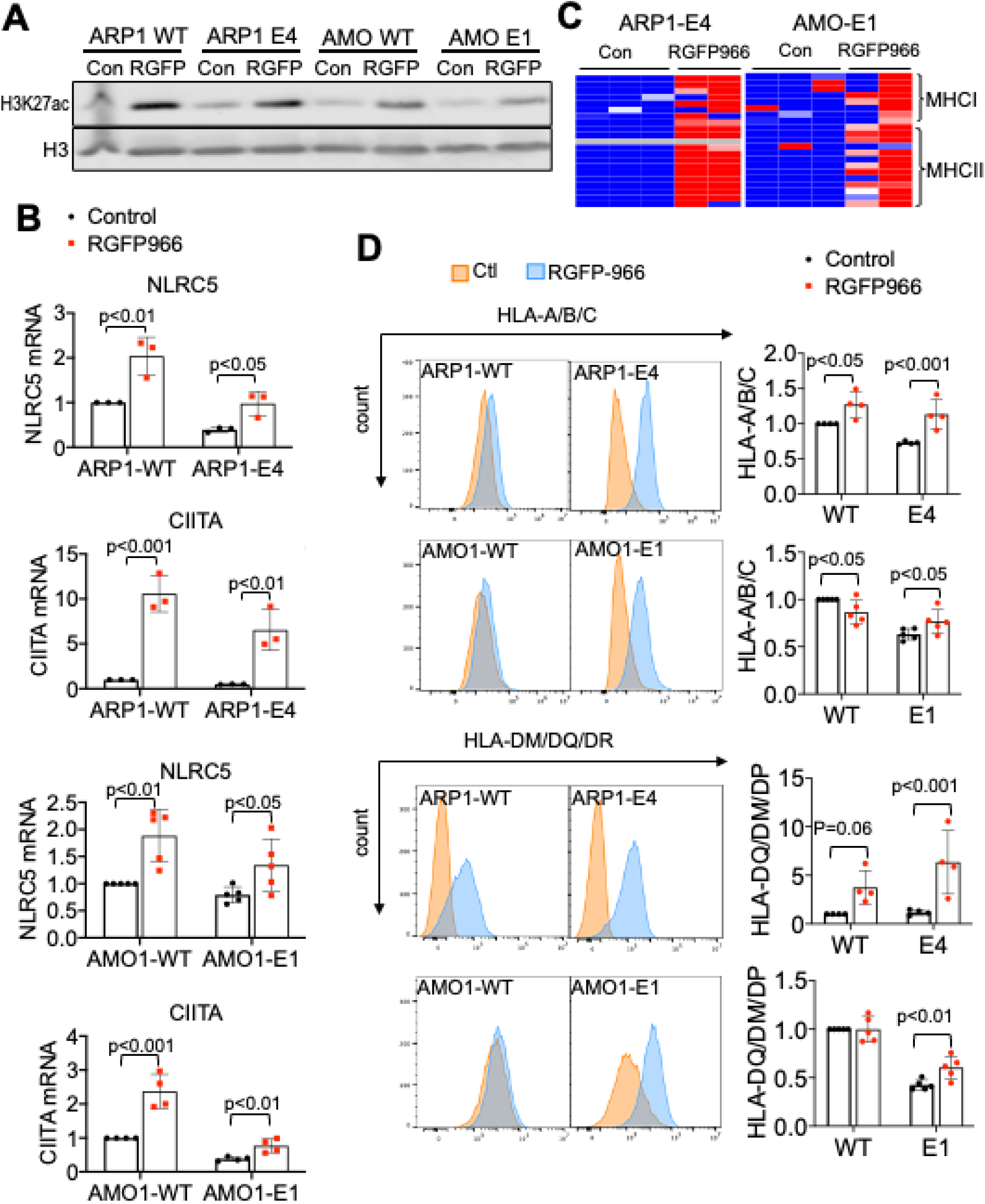
HDAC3 inhibition restores MHCs expression in MM cell lines. **A**) Immunoblot of H3K27ac 24h post treatment with 10µM RGFP966. **B)** Heatmap of z-scores for all expressed MHC genes in KDM6A clonal isogenic cell lines. **C)** mRNA analysis by QPCR normalized to GAPDH 24h post treatment with 10µM RGFP966 (3 to 5 biological replicates; +/− SD. Wilcoxon T-test). **D,** Flow cytometry analysis of HLA-A/B/C and HLA-DM/DQ/DR in myeloma isogenic cell lines treated or not with 10µM RGFP966 for 24h or 72h. Panel on the right show the histograms MFI quantification of HLA-A/B/C and HLA-DM/DQ/DR (4 or 5 biological replicates; +/− SD, Wilcoxon T-test).

### KDM6A depletion decreases tumor immunogenicity *in vivo*

To test whether decreased MHCs expression caused by KDM6A loss may affect tumor immunogenicity *in vivo*, we developed a mouse embryonic fibroblast (MEF) cell line by 3T3 immortalization protocol (32) from C57JBL/6J mice that possess loxP sites flanking exon 3 of *Kdm6a* (33). These fibroblasts were infected with an adenovirus expressing Cre recombinase to disrupt *Kdm6a* or a control adenovirus harboring GFP (**Fig. 6A**). Because activating mutations of the Ras/MAPK pathway are found in about 50% of MM (34),we transformed wild type and *Kdm6a* mutant cells by transducing a retrovirus expressing KRAS harboring the G12V mutation. There was modestly increased growth in *Kdm6a* KO MEF (Supplementary Fig. 8A). Growth was increased in KRAS expressing cells but did not differ between KDM6A null and replete cells (Supplementary Fig. 8A). RNA-seq showed that expression of immune genes was also affected in *Kdm6a* knockout fibroblasts compared to GFP controls. All MHC class I genes expressed in wild type fibroblasts harboring oncogenic RAS were downregulated in the knockout cells (**Fig. 6B**) along with *Nlrc5* (**Fig. 6C**) and genes involved in antigen presentation and Type I interferon signaling (Supplementary Fig. 8B). Flow cytometry confirmed downregulation of MHC-I on the cell surface of *Kdm6a* null cells **(Fig. 6D**). As in the case of the myeloma cell lines, loss of KDM6A was associated with decreased in H3K27 acetylation at the *Nlrc5* locus (Supplementary Fig. 8C). HDAC3 inhibition by RGFP-966 increased H3K27 acetylation (Supplementary Fig. 8D), Nlrc5 mRNA levels (Supplementary Fig. 8E) and partially restore the expression of MHC class I in *KDM6A* null MEFs (Supplementary Fig. 8F).

**Figure 6:**
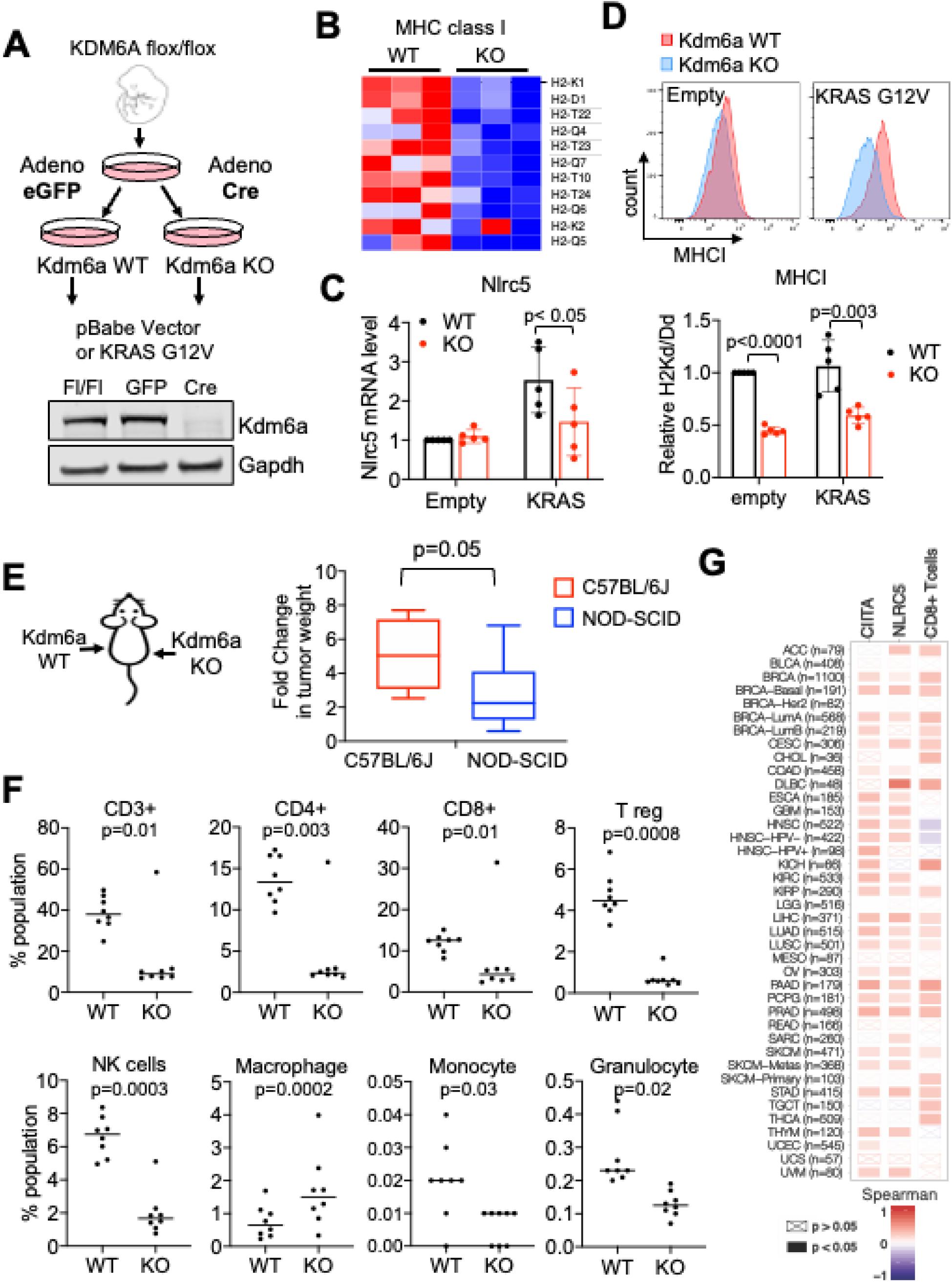
KDM6A depletion decreases MhcI expression in MEF and decrease tumor immunogenicity *in vivo*. **A**) Schematic of the protocol used to develop isogenic *Kdm6a* knockout murine embryo fibroblasts (MEF) from a C57BL/6J mouse homozygous for an allele of *Kdm6a* in which exon 3 was flanked with LoxP sites. **B)** Heatmap of z-score for all expressed MHC-I genes in *Kdm6a* wild type and knockout MEF. **C)** *Nlrc5* mRNA analysis by QPCR normalized to GAPDH (5 biological replicates; +/− SD. Mann Whitney T-test) **D) Upper panel:** Flow cytometry analysis of H-2kb in *Kdm6a* wild type or knockout MEFs. **Lower panel:** MFI quantification of H-2kb surface expression from upper panel (5 biological replicates; +/− SD, Mann Whitney T-test). **E)** RAS-transformed wild type and *Kdm6a* knockout MEFs were injected into left and right flanks respectively of C57BL/6J and NOD-SCID mice and tumors excised and weighed after 3 weeks. The ratio of tumor weights from animal injected with *Kdm6a* replete or *Kdm6a* deficient Ras transformed fibroblasts was calculated in immunocompetent C57BL/6J and immunodeficient NOD-SCID mice. (8 biological replicates; +/− SD, Mann Whitney T-test). **F)** Flow cytometry quantifications of T cell, NK cells and macrophage populations in the tumors isolated in **E**. **G)** Spearman correlation between KDM6A and NLRC5 or CIITA gene expression and CD8+T cells infiltration in tumors isolated from various human cancers and correlation between KDM6A expression and T cell infiltrates as determined using Cistrome Timer 2.0 (35).

KRAS transformed *Kdm6a* replete or *Kdm6a* deleted fibroblasts were injected into each flank of immunocompromised NOD-SCID mice and immunocompetent C57BL/6 mice. In immunocompromised mice tumors formed on both flanks, and *Kdm6a* null tumors were twice the size of *Kdm6a* wild-type masses. By contrast, in immunocompetent mice *Kdm6a* null tumors grew 5 times the size of *Kdm6a* wild-type tumors (**Fig. 6E)**. Together these observations suggest that tumor suppressive functions of KDM6A are partially dependent on an intact immune system. Flow cytometry analysis of immune cell infiltrates of *Kdm6a* replete tumors grown in C57JBL/6J mice indicate an active immune environment. By contrast, *Kdm6a* knockout tumors exhibited a marked decrease in infiltrating T cells (CD4+, CD8+ and Treg), NK cells, granulocytes and monocytes (**Fig. 6F**) but an increase in macrophage inflitration. Using the Cistrome Timer 2.0 database (35), we found a positive correlation between *KDM6A* and both *NLRC5* and *CIITA* expression in a variety of tumor types, as well as a positive correlation between *KDM6A* expression and CD8 T cell infiltration within the tumor (**Fig. 6G),** supporting an important role of *KDM6A* in the immune response of a variety of tumors. Collectively, these data suggest that *Kdm6a* suppresses growth in part by supporting tumor immunogenicity by maintaining expression of MHCI, MHC-II, and critical cytokines.

## Discussion

KDM6A is a regulator of enhancer activity and a tumor suppressor in MM. Here, we found that KDM6A binds to enhancers associated with genes regulating immune recognition pathways such as antigen presentation, chemokine secretion and T cell trafficking to tumors. KDM6A binds and regulates transcription of the master regulators of MHC-I and MHC-II, NLRC5 and CIITA. KDM6A loss was associated with a marked decreased in H3K27ac at active enhancers while H3K27me3 was increased in the bodies of inactive genes in the proximity of KDM6A. Restoration of histone acetylation using a HDAC3-selective inhibitor upregulates MHC-I and II expression in KDM6A deficient MM cells. We further found that loss of KDM6A compromised the immune response to a K-Ras driven malignancy, relevant to MM given that nearly 50% of MM patients have activating mutations of the Ras/MAPK pathway (36,37).

We found that both active and poised enhancers bound by KDM6A are associated with genes related to immunity. Consistent with this, we observed diminished upregulation of IFNα target genes upon stimulation by IFNα in KDM6A depleted cells. Analysis of chromatin at KDM6A bound enhancers revealed its ability to modulate histone acetylation at H3K27 but surprisingly, not H3K27 methylation. This is consistent with data from Wang et al., where H3K27me3 levels at enhancers bounded by KDM6A are not affected by its depletion (29).

We found that methylation of H3 at lysine K27 increased in KDM6A null cells, specifically in the body of genes in close proximity to enhancers and promoters normally bound by KDM6A. This contrast with observations from other studies of KDM6A that focused on enhancers specific effects or relatively short genomic areas (28). Whether increased H3K27me3 is a cause or an effect of decreased gene expression is unknown. Lack of transcription was previously shown to trigger H3K27me3 accumulation in gene bodies (38). Thus, KDM6A loss may cause decreased gene expression through a decrease in H3K27ac which in turn leads to H3K27me3 accrual. Our findings suggest that KDM6A has a dual function regulating distal H3K27ac at enhancers and concurrently controlling proximal methylation of H3K27 in the body of genes covering a wide genomic area. Disrupting KDM6A enzymatic activity did not reduce H3K27ac levels to that observed in KDM6A null cells, suggesting a structural or scaffolding function of KDM6A to facilitate the recruitment or activity of CREBBP or EP300 and the COMPASS-complex at enhancers.

KDM6A, along with KMT2C/D and SWI/SNF components are collectively critical for activation of enhancers though COMPASS-like complexes. KDM6A localization across the genome in MM cells closely overlaps with previously defined patterns of EP300 as well as CREBBP, consistent with ability to interact with these histone acetyl transferases (reviewed in (39)). KDM6A appears essential for the integrity of the COMPASS-like complex and in its absence, there is decreased accessibility and acetylation of immune response gene enhancers (29). KDM6A binding sites are enriched for motifs of IRF, STAT and E2F transcription factors that were found to interact with KDM6A, CEBPB and EP300 (BIOGRID database) which could be responsible for the recruitment of these complexes to key enhancers, facilitating activation of genes implicated in immune recognition of tumors. Although generally recognized as a transcriptional repressor, IRF2 was found to act as a transcriptional activator for many key components of the MHC-I pathway, including the immunoproteasome, TAP, and ERAP1. Furthermore, IRF2 is frequently downregulated in cancer (40).

KDM6A loss led to decreased MHC-I and MHC-II gene expression in MM cell lines as well as murine fibroblasts. Loss of MHC-I expression or loss of the capacity to induce upregulation of MHC-I cell surface expression commonly occurs in malignant cells and serves as a strategy to escape T-cell recognition (41). Moreover, expression of NLRC5 by cancer cells was shown to enhance cytotoxic cytokine production and cancer cell killing by CD8+ T cell. Prior studies demonstrated that KDM6A deficiency decreased immunogenicity of bladder cancer (42) by promoting cytokine/inflammatory pathways (43). In addition, KDM6A expression has been identified as a marker for response to vaccine therapy in bladder cancer (44), suggesting that KDM6A supports immune responses. A study in a medulloblastoma mouse model revealed that T cell attraction to tumors was promoted through a KDM6A-dependant activation of Th1-type chemokine expression (45). Consistent with this, we found that secretion of inflammatory and chemotactic cytokine for T-cells was inhibited by the loss of KDM6A and that *Kdm6a* null Ras-driven tumors displayed decreased T cell infiltration.

A growing list of tumor-associated mutations has been described which mediate immune escape by multiple mechanisms. Notably, a recent screen in which murine tumor cells were gene edited and then grown *in vivo* in both immunocompetent and immunodeficient mice found an enrichment for the loss of tumor suppressor genes in the presence of an adaptive immune system. Interestingly, an important subset of these tumor suppressor genes encoded chromatin modifiers including KDM6A (46). Other studies support the idea that mutations or deregulation of epigenetic modifiers are associated with immune escape of tumors (47). For example, in several malignancies PRC2, which opposes the action of KDM6A to promote gene silencing, is elevated and represses expression of MHC class I antigen presenting (48–50). We extended these findings by showing that KDM6A also binds enhancers for CIITA and NLRC5 activating their transcription which in turn stimulate expression of MHC-I and MHC-II gene respectively. Loss of MHC-I expression will have a major effect on presentation of tumor neoantigen to T-cells. MHC-II and CITA loss could also modulate antitumor immunity in myeloma. MHC-II levels drop as normal B cells differentiate into plasma cells, but multiple myeloma cells can express MHC-II and act as antigen presenting cells. Increasing evidence shows that the MHC-II antigen presenting-complex is expressed in tumors of various tissue origin despite being typically associated with professional antigen-presenting cells. MHC-II expression in tumors is associated with better outcomes in patients with cancer and with tumor rejection in murine models (Reviewed in). Therapeutic MHC-II upregulation is thus considered as a possible treatment maneuver in cancer and our data supports considering this strategy in MM.

Activation of NF-κB and p300/CREBBP was shown to potentiate cancer chemoimmunotherapy through induction of MHC-I antigen presentation (55). We further show that enhancing histone acetylation using HDAC3 inhibitors helps to restore the expression of NLRC5, CIITA and their respective MHC targets. HDAC3 inhibition also increased MHC expression in *Kdm6a* replete cells suggesting this agent might have a beneficial effect in multiple situations in which MHC expression is depressed in cancer cells.

KDM6A loss and its ability to help cells escape immune surveillance may apply to other cancers. Inactivating mutations of *KDM6A* as well as *KMT2C, KMT2D* and *SWI/SNF* components of its complex are among the most common found in all human cancers (56). For example, *KDM6A* is mutated (57) in Breast cancer (58), Clear Cell Renal Cell Carcinoma (CCRCC) (59), Lung cancer, Medulloblastoma, bladder carcinoma (60), acute lymphoblastic leukemia (61) and chronic myelomonocytic leukemia. In addition, the loss of X chromosome and X-linked gene expression occurs in 22% of all cancers. As such, pharmacological interventions to restore MHC and other immune function in tumor cells that have lost the activity of the enhancer activity complexes should be considered as part of the treatment approach to KDM6A mutant MM and other malignancies.

## Materials and Methods

### Cell lines

The multiple myeloma cell line ARP-1 RRID:CVCL_D523 was established at the University of Arkansas for Medical Sciences (62). The RPMI-8226 RRID:CVCL_0014, L363 RRID:CVCL_L363, EJM RRID:CVCL_2030, KMS12 RRID:CVCL_1334, AMO-1 RRID:CVCL_1806 and Karpas-620 RRID:CVCL_1823 cell lines were gifts of the late Michael Kuehl, National Cancer Institute. Cells were maintained in advanced RPMI-1640 medium (Invitrogen, Frederick, USA), supplemented with 5% fetal bovine serum (FBS), or 20% FBS (Karpas-620); or IMDM medium containing 10% FBS (EJM). All growth medium were supplemented with 1% penicillin (100 units/ml), and 1% streptomycin (100 μg/ml). Cell line authenticity was confirmed by short tandem repeat (STR) profiling (Lab Corp Genetica, Burlington, NC).

### Cell Proliferation Assay

Equal volume of CellTiter-Glo Reagent to the volume of cell culture medium were added into each well and mixed for 2 minutes on a shaker to induce cell lysis. The plates were incubated at room temperature for 10 minutes and luminescence was measured with the plate reader CLARIOstar (BMG LABTECH).

### Carboxyfluorescein succinimidyl ester (CFSE) proliferation assay

Cells were labeled with 2 μM CFSE (Selleckchem, Houston, USA) at 37 °C for 20 min. Proliferation was determined by detecting the fluorescence intensity of CFSE via flow cytometry using Beckman CytoFLEX (Indianapolis, Indiana) and analysed using FlowJo v10 software.

### CRISPR-Cas9 gene editing

Annealed guide RNA/tracer RNA ribonucleotides were mixed with Cas9 protein (IDT, Coralville) at a ratio of 1:1 (0.8μg of each) at room temperature for 5 min. The complex or Cas9 only (control wild type cells) was mixed with a non-specific oligonucleotide (IDT) and electroporated into 5×10^5^ ARP1 cells using a NEON (ThermoFisher Scientific, Waltham, MA) apparatus at a setting of a 20 msec pulse at 1600 Volts. The guide RNA recognition sites are the following: *KDM6A* exon 1: TGCGTTTCCATGAAATCCTG; *KDM6A* exon 4 (clone E4): CAGCATTATCTGCATACCAG; KDM6A exon 6 (clone E6): AGCTTTTGTCGAGCCAAGGA. Two days post-electroporation, clones were isolated by limiting dilution. Identification of KDM6A mutants was performed by Sanger sequencing of the amplified targeted region. Primers for generation of amplicons were: *KDM6A* exon 1 forward; 5’-CATGAAATCCTGCGGAGTGT-3’, reverse; 5’-TACTCGTTAACGCTCAGGGA-3’. *KDM6A* exon 4 forward; 5’-TGT GGT GGG AAT CTT GTT ACC-3’, reverse; 5’-GCA CAA ACA TAA ATA CTC TCA ACC C-3’. *KDM6A* exon 6 forward; 5’-GTT TCA ATG TAC TAC CAA GCA AGA A-3’, reverse; 5’-ACC CAA CAA CCT ACC TTT AAA CT-3’. Deep sequencing of the amplicons was performed at the Center for Computational and Integrative Biology at Massachusetts General Hospital (Boston).

### Lentiviral transduction

ARD, L363 and KMS12 cells were transduced with the lentiviral vector pInducer20 (RRID:Addgene 44012, Gift of Stephen Elledge) harboring a Tet operator-controlled KDM6A cDNA (pKDM6A) or a control plasmid (pLac). Lentiviruses were generated by transfection of 293T cells with these plasmids, (10) in addition to the packaging vectors psPAX2 (RRID:Addgene_12260) and pMD2.G (RRID:Addgene_12259) gifts of Didier Trono (63), using Lipofectamine P3000 (ThermoFisher Scientific). Viral supernatant was collected, passed through 0.45um filter and concentrated with LentiX Concentrator (Takara Bio USA, Mountainview, CA). Concentrated lentivirus was added to cells in the presence of 6 μg/ml polybrene (MilliporeSigma, Burlington, MA) and centrifuged at 1000xg for 90 minutes. Selection with 2 mg/ml G418 was initiated 48h post transduction. For induction of KDM6A, cells were grown in the presence of 200 ng/ml doxycycline for 6 days. Retrovirus were generated by transfection of 293T cells with these plasmids, using Lipofectamine P3000 (ThermoFisher Scientific).

### Immunoblotting

Nuclear proteins were extracted using the Nuclear Complex Co-IP Kit (Active Motif, Carlsbad, CA), following the manufacturer’s instructions. For total protein extraction, cells were disrupted in 1% NP-40 lysis buffer (140 mM NaCl, 10 mM Tris-HCl pH 8, 1% NP-40) supplemented with proteinase inhibitors (Roche). Proteins were separated by electrophoresis (SDS-PAGE), blotted, and detected using the Odyssey^®^ CLX imager, LI-COR (Bioscience). Primary antibodies were KDM6A (clone DQ1I) Cell Signaling (Danvers, MA) #33510 RRID:AB_2721244 and HDAC2 (clone 3F3) Millipore Sigma #05-814 RRID:AB_310022. Secondary antibodies used are IRDye^®^ 800CW Goat Anti-Mouse and IRDye® 680RD Goat anti-Rabbit (LI-COR Bioscience). All primary antibodies were used at 1:1000 dilution and secondary antibodies at 1:10000 in TBS with 0.1% Tween-20.

### Analysis of cytokines and chemokines

Cytokines were measured using a Human Magnetic Luminex Assay (Bio-Techne, Minneapolis, MN, USA), a magnetic bead-based sandwich immunoassay for cytokines, in accordance with the manufacturer’s instructions. Briefly, 96-well filter bottom plates are wetted with buffer, and beads conjugated with capture antibodies for each cytokine are added. Serial dilutions of cytokine standards and quality controls are designated by the manufacturer’s recommendation and followed by the addition of 25 µl of lysate in triplicates. Plates are washed on mixer for 30 min at room temperature then incubated 18 hours at 4°C. The plates are washed, then incubated with a biotin-labeled human cytokine detection antibodies for one hour at room temperature with shaking. Plates are incubated with streptavidin-PE for 30 minutes, then final set of washes are performed and the beads resuspended in washing buffer. The beads were analyzed in a Luminex FlexMap3D instrument (Luminex Corporation, Austin, TX) to record the median levels of fluorescent per bead population.

### Mouse embryonic fibroblasts

*Kdm6a^flox/flox^* mice were generated from the JM8.N4 ES cell line obtained from EUCOMM by the laboratory of Timothy Ley and Lucas Wartman (Washington University, St. Louis MO). The neomycin cassette in the embryonic stem cell line was removed by FLP recombinase prior to development of the mouse. These mice have a floxed exon 3 of the Kdm6a gene. Deletion of the third *Kdm6a* exon produces a frameshift and introduction of a translational stop codon when *Kdm6a* is spliced from exon 2 to exon 4. MEFs were isolated from embryonic day 12.5 *Kdm6a^flox/flox^* mice by uterine dissection for individual embryos. Embryos were rinsed with PBS, followed by removal of the mouse head and embryonic internal organs. The embryo body was transferred to a clean culture dish containing 3 ml of 0.25% trypsin-EDTA and then forced through a 1-ml syringe with a 16-gauge needle. The tissue homogenate was incubated for 30 min at 37 °C in 10 ml trypsin-EDTA solution, and cell suspension was divided into two or three 10-cm tissue culture dishes in DMEM. Cells were maintained in DMEM medium containing 10% FBS supplemented with 1% penicillin (100 units/ml), and 1% streptomycin (100 μg/ml). MEFs were immortalized after 15 passages by the 3T3 protocol. Cre recombinase was delivered to *Kdm6A^flox/flox^* MEFs by adenoviral transduction of the Ad5CMVCre-eGFP or control Ad5CMVeGFP virus (University of Iowa Viral vector Core). Cre mediated disruption of *Kdm6a* was confirmed by genotyping as previously described (64). MEF cells were transduced with the retroviral vector pBABE-puro-KRAS^G12V^ (Gift from Christopher Counter (RRID:Addgene 46746) or a control plasmid pBABE-puro (Gift from Hartmut Land, Jay Morgenstern and Bob Weinberg (RRID:Addgene_1764)). Retrovirus was generated by transfection of 293T cells with these plasmids, using Lipofectamine P3000 (ThermoFisher Scientific). Viral supernatant was collected, passed through 0.45µm filter and concentrated with RetroX Concentrator (Takara Bio USA, Mountainview, CA). Concentrated lentivirus was added to cells in the presence of 6 μg/ml polybrene (MilliporeSigma, Burlington, MA) and centrifuged at 1000xg for 90 minutes.

### Flow cytometry

Myeloma cell lines (2× 10^5^ –1×10^6^ cells) were washed in PBS and then labeled with 1 µl of a mix of antibodies as indicated in 50 μl PBS. Staining was performed at 4 °C for 15– 30 min, followed by washing with 1 ml of PBS before the analysis on the CytoFLEX LX (Beckman Coulter, Indianapolis). Antibodies used are the following: HLA-DR, DP, DQ clone Tü39 (Biolegend #361716), HLA-A/B/C clone W6/32 (Biolegend #311413), mouse MHC-I FITC conjugated H2-Dd/H-2Kd clone 34-1-2S (Invitrogen #11-5998-82), mouse MHC-I PE conjugated H-2Kb clone AF6-88.5.5.3 (Invitrogen #12-5958-82).

### Quantitative real-time PCR

Total RNA was extracted using TRIzol reagent (ThermoFisher Scientific) and cDNA was synthesized from 1μg RNA using iSCRIPT as per the manufacturer’s instructions (Biorad, Hercules, CA). qPCR analysis of samples was performed on the CFX96 Real-Time System (Biorad) with Sso Advanced Universal SYBRGreen supermix (Biorad) with the following thermal cycler conditions for all reactions: 1 cycle 95 °C for 10 min, followed by 40 cycles at 95 °C for 10 s, and at 60 °C for 30 s, ending with 72 °C incubation for 5 min. For cDNA qPCR data was analyzed using the ^ΔΔ^Ct method (65) normalizing to GAPDH as the housekeeping gene. qPCR primer sequences are as follows:

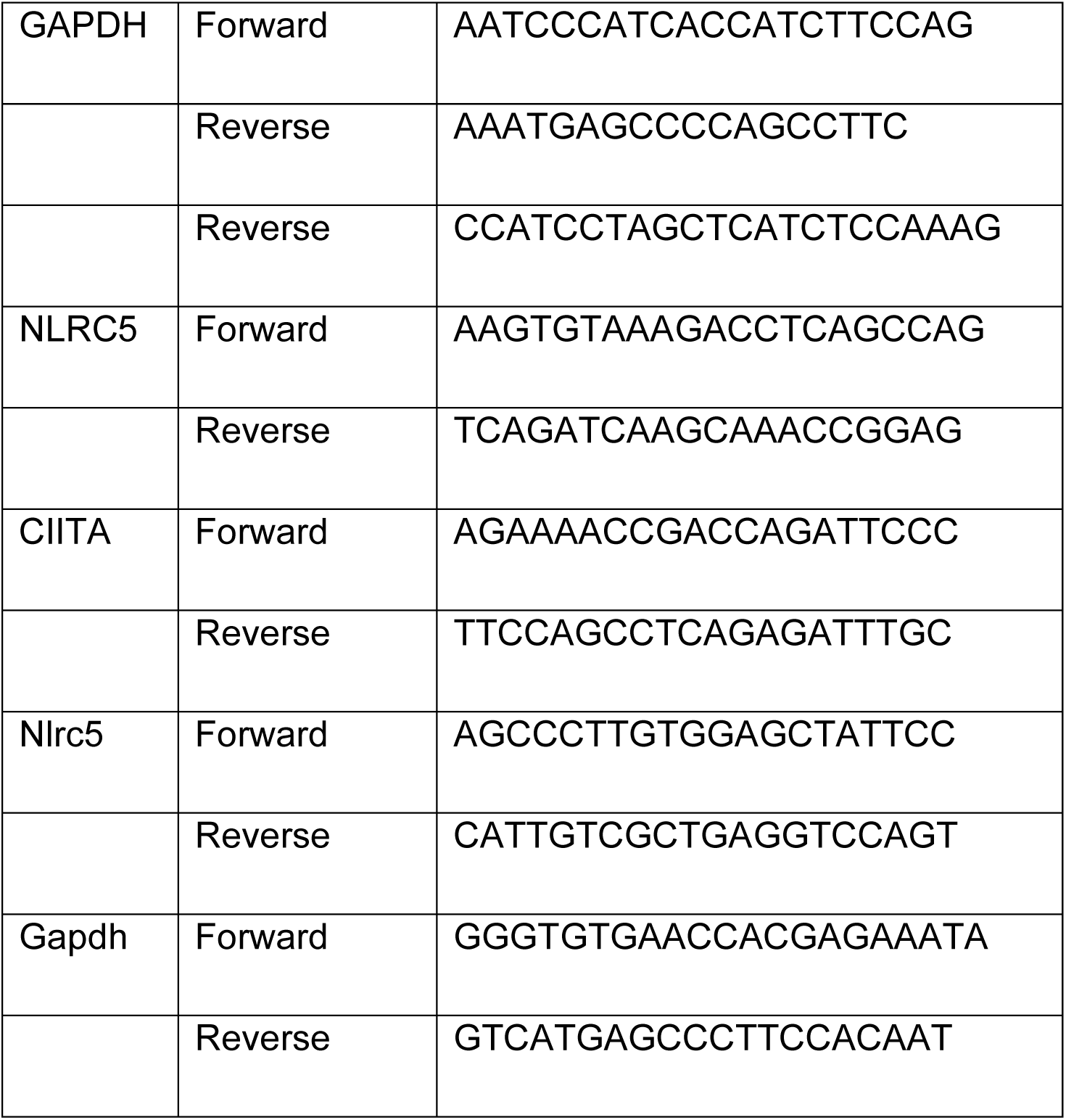

### Mouse Xenografts

MEFs were resuspended in Matrigel (Collaborative Research) diluted 1:5 with DMEM media at a concentration of 10×10^6^ cells per ml. Each 6–9 week-old mouse was injected with a 200µl of Matrigel cell suspension of 0.33× 10^6^ cells per flank of RAS transduced *Kdm6a* wild type or knockout cells. When tumors size reached the endpoint of 2 cm diameter (3 weeks post-transplant), mice were euthanized and tumors excised, weighted and 200mg was processed for flow cytometric analysis of immune infiltrate. Tumors were detached from the mice and approximately 150 mg of tumor tissue was enzymatically and mechanically digested using the Mouse Tumor Dissociation Kit (Miltenyi Biotec) to obtain a single cell suspension. Red blood cells were removed from single cell suspension using ACK lysis buffer and mononuclear cells were isolated by density gradient using SepMate Tubes (StemCell Technologies) and Lymphoprep density gradient media (StemCell Technologies). Subsequently, the cells were incubated with combinations of the following antibodies: anti-mouse CD62L-BV785 (clone MEL-14, RRID:AB_398533), anti-mouse MHC-II I-A/I-E-BB515 (Clone 2G9, BD Biosciences, 1:400), anti-mouse CD11B-PEdazzle (clone M1/70, 1:200), anti-mouse CD45-AF532 (clone 30F.11, RRID:AB_11218871), anti-mouse CD3-APC-Cy7 (clone 17A2, RRID:AB_1727461), anti-mouse CD8-BV510 (clone 53-6.7), anti-mouse CD4-BV605 (clone GK1.5, RRID:AB_2564591), anti-mouse NK1.1-AF700 (clone PK136, RRID:AB_2574504), anti-mouse CD69-SB436 (clone H1.2F3, eBioscience, RRID:AB_2688109), anti-mouse CD279 (clone-PerCP-EF710, eBioscience Inc), anti-mouse CD366-AF647 (clone B8.2C12, RRID:AB_1626175), anti-mouse CD11C-PE-Cy7 (clone N418, RRID:AB_469589), anti-mouse Ly6G-APC (clone IA8, RRID:AB_2573307), anti-mouse Ly6C-BV711 (clone HK1.4, RRID:AB_2562630), anti-mouse F4/80-BV650 (clone BM8, RRID:AB_2925696), anti-mouse CD80-BV480 (clone 16-10A1, RRID:AB_465132, BD Biosciences), anti-mouse CD25-PE-Cy5 (clone PC61) plus FVD-EF780 (eBioscience) and mouse FcR blocker (anti-mouse CD16/CD32, clone 2.4G2, RRID:AB_626927, BD Biosciences). Next, cells were fixed and permeabilized using the FOXP3/Transcription Factor Staining Buffer Set (eBioscience). After fixation, cells were stained with a combination of the following antibodies: anti-mouse FOXP3-eFluor 450 (clone FJK16S, 1:50, eBioscience, RRID:AB_1518813), anti-mouse Granzyme B-FITC (clone GB11, RRID:AB_1645488), anti-mouse Perforin-PE (clone S16009B, RRID:AB_2721641), anti-mouse Ki-67-PerCP-Cy5.5 (clone 16A8, RRID:AB_2629530). Flow cytometry was performed on a 3 laser Cytek Aurora Cytometer (Cytek Biosciences, Fremont, CA) and the data was analyzed using FlowJo software (BD Biosciences, RRID:SCR_008520). All listed antibodies for flow cytometry are from Biolegend, unless otherwise specified and were used at 1:100 dilution for this experiment.

### Quantification and Statistical Analysis

Additional statistical analysis, not specific to a particular protocol were carried out using GraphPad Prism 8 (RRID:SCR_002798). In general, data are reported as mean ± SD. Independent biological replicates are shown as individual data points, as indicated in the figure legends.

### Financial support

Research reported in this publication was supported by R01CA180475, the Leukemia and Lymphoma Society Specialized Center of Excellence, Samuel Waxman Cancer research Foundation (JDL), Leukemia and Lymphoma Society Special Fellow Award (DDR and JL), the National Cancer Institute of the National Institutes of Health under award number K22CA266739 (BGB), CA260239 (WZ) and CA269661 (WZ), the Paula and Roger Riney Foundation (LHB, BGB), Rally Foundation for Childhood Cancer Research and Bear Necessities Pediatric Cancer Foundation (JL), Investigador AECC award from the Fundación AECC INVES19059EZPO (TE).

Table S1: KDM6A mutants allele frequency detected by amplicon sequencing.

Table S2: KDM6A binding site and associated genes in ARP1 cell lines and in Karpas-620.

Table S3: RNA expression in myeloma isogenic clonal cell lines.

Table S4: Motif enrichment analysis obtained with HOMER.

