## Supplementa figures for "KDM6A Regulates Immune Response Genes in Multiple Myeloma"

Figure S1

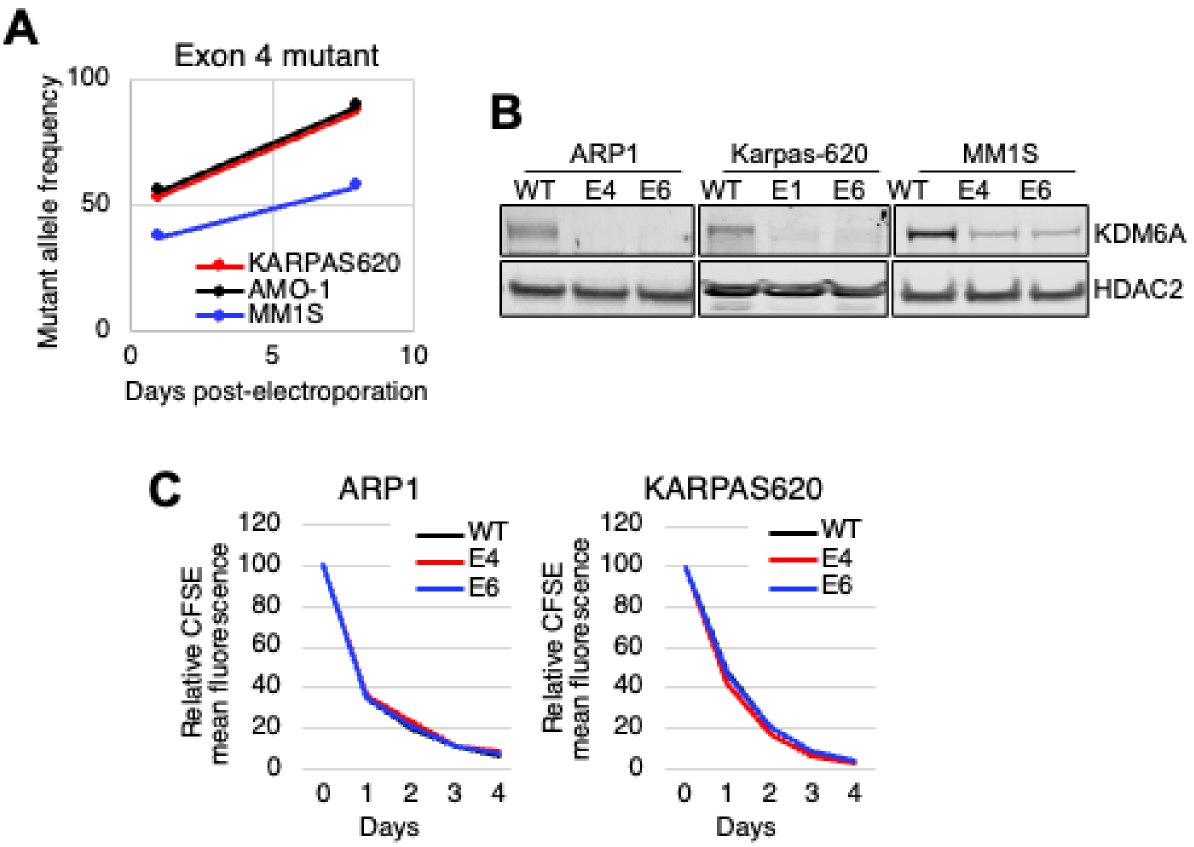


**Figure S1: KDM6A loss provides a transient proliferative advantage in MM cells. A)** Mutant allele frequency was measure by deep-sequencing 1 and 8 days after electroporation with RNP containing Cas9 and gRNA directed against KDM6A exon 4 in MM1S, KARPAS-620 and ARP1 cell lines. **B)** Immunoblot showing downregulation of KDM6A protein in CRISPR edited cell pools two weeks after electroporation with RNPs containing Cas9 and gRNA directed against exon 4 (E4) and 6 (E6) of the *KDM6A* locus (KARPAS, MM1S, ARP1). **C)** KDM6A wild type and knockout cells were stain by carboxyfluorescein succinimidyl ester (CFSE) and fluorescence was detected at the time points indicated.

Figure S2

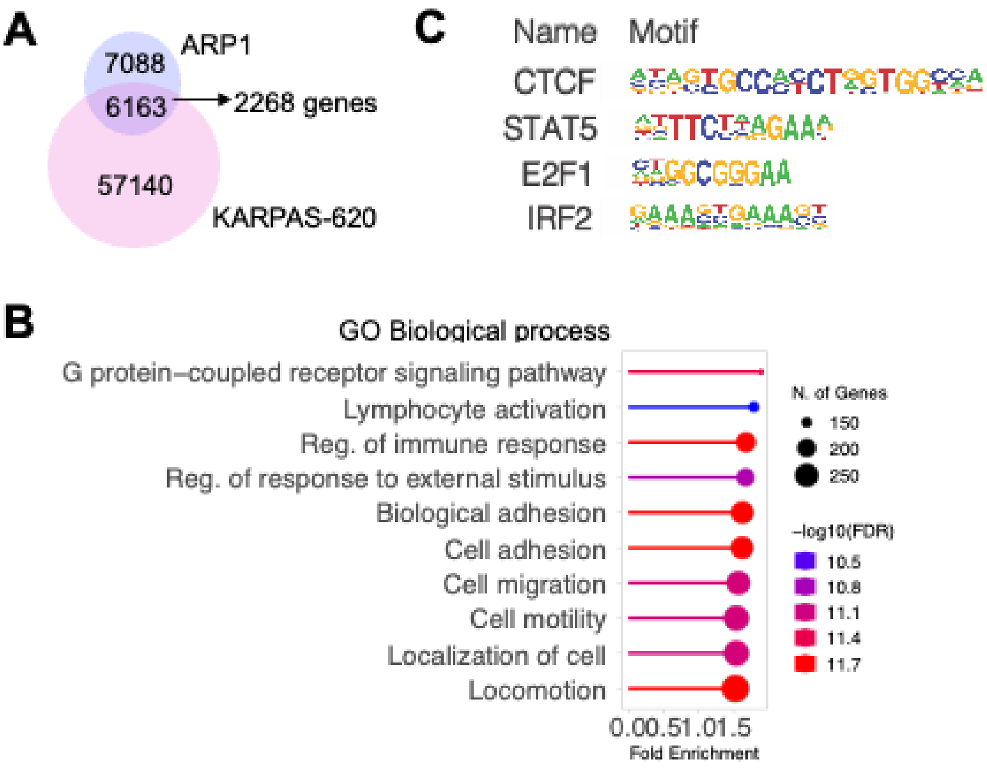


**Figure S2: KDM6A binding sites in Myeloma cell lines**. **A)** Overlap of KDM6A binding sites detected in ARP-1 and KARPAS-680 cell lines. **B,** Dot plot visualization of GO biological process enrichment analyses of genes found within 100kb from KDM6A KARPAS620 cell line using the GoShiny v0.77 tool. The size of the dots scale with the number of genes and the color scale with FDR. **C**) HOMER-identified enriched transcription factor binding motif in KDM6A binding site found commonly in KARPAS-620 and ARP1 cell lines.

Figure S3

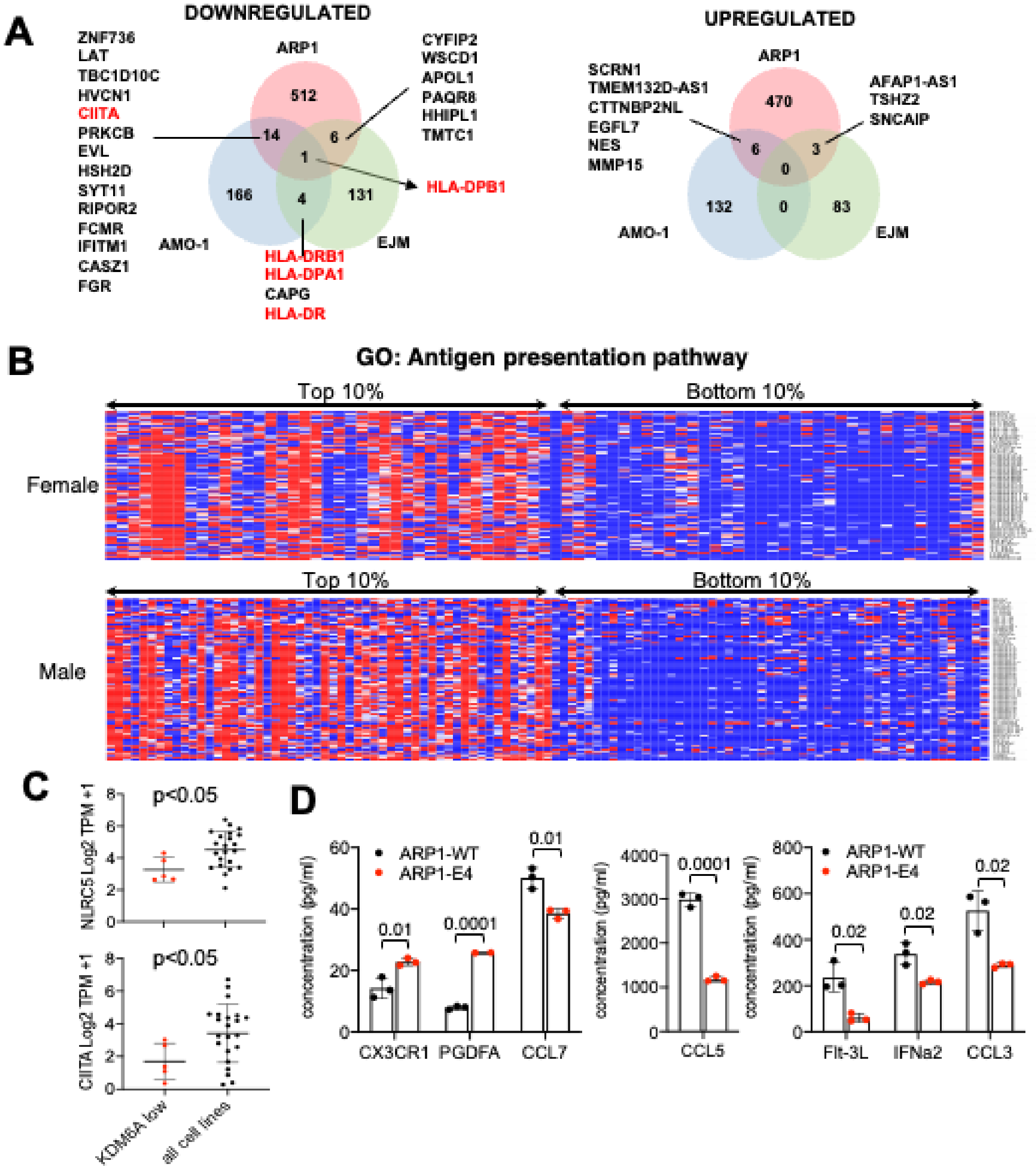


**Figure S3: KDM6A controls expression of genes involved in cell immunity. A)** Venn diagram showing overlap of genes downregulated and upregulated (1.5 fold change, $p < 0.05$) in KDM6A knockout compared to WT clonal isogenic cell lines ARP1, AMO-1 and EJM. **B)** Heatmap of z-scores for gene included in the annotated GO: Antigen presentation pathway in the top expressing highest and lowest KDM6A. **C)** Comparison of NLRC5 and CIITA gene expression (TPM) with KDM6A gene expression in myeloma cell lines using Depmap data (Mann-Whitney T-test). **D)** Multiplex ELISA analysis of cytokine in ARP-1 WT and KO clonal cells line (Mann-Whitney T-test).

Figure S4

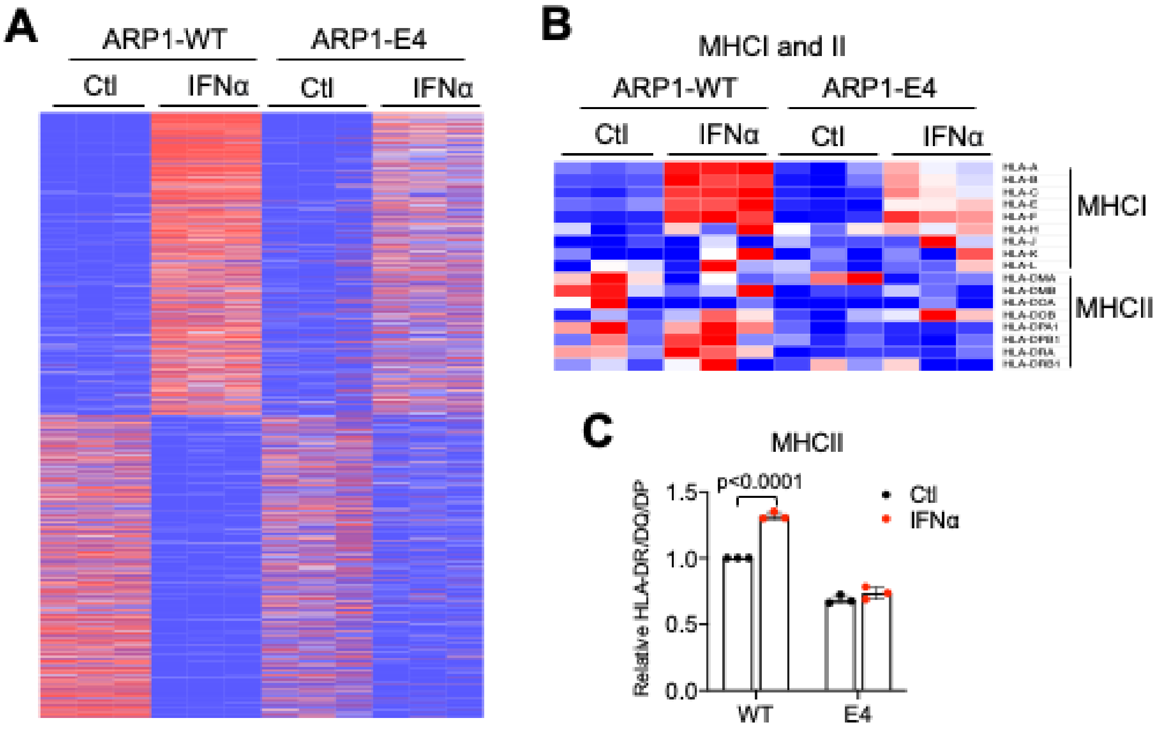


**Figure S4: Interferon alpha response is attenuated in KDM6A KO cells. A,** Heatmap of gene expression (TPM) for gene significantly deregulated by treatment with 200ng/ml IFNα for 18h. **B,** Heatmap of gene expression (TPM) of all expressed MHC genes +/- IFNα. **C,** Flow cytometry quantification (MFI) of HLA-A/B/C and HLA-DM/DQ/DR in clonal isogenic cell lines +/- IFNα.

Figure S5

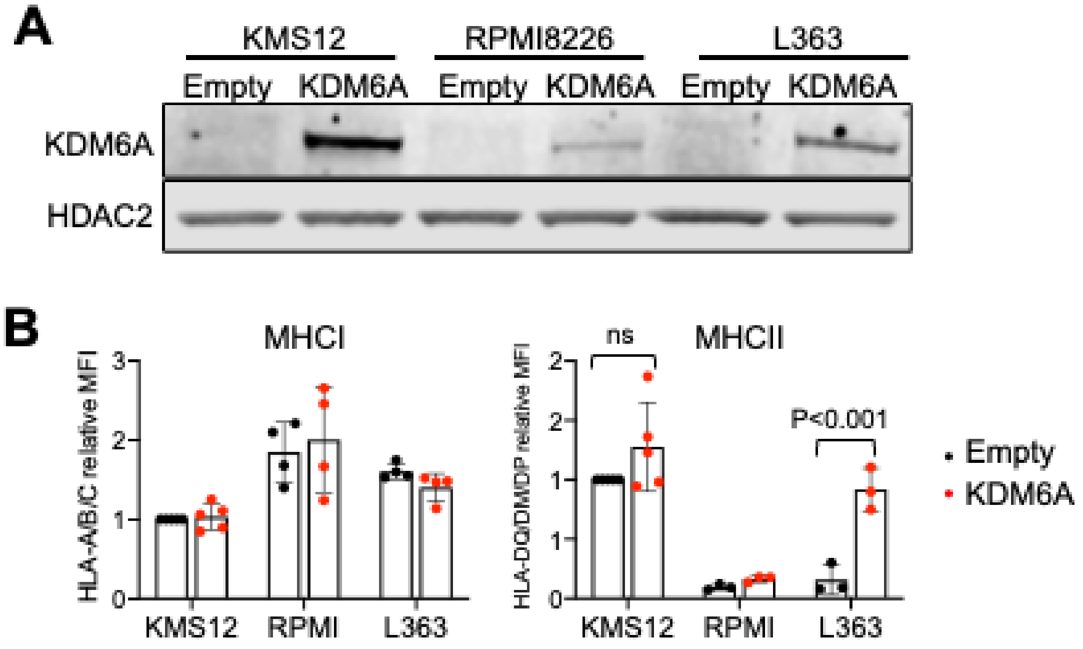


**Figure S5: Stable add-back of KDM6A can induce MHCII expression. A** Immunoblot analysis showing induction of KDM6A protein expression after 3 days treatment with 1ug/ml doxycycline in stable cell lines expressing inducible KDM6A **B,** Flow cytometry quantification (MFI) quantification of HLA-A/B/C and HLA-DM/DQ/DR surface expression in KDM6A inducible cell lines L363, RPMI8226 and KMS12, 6 days under treatment with 200ng/ml doxycycline. (3 or more biological replicates; +/- SD. Wilcoxon T-test ).

Figure S6

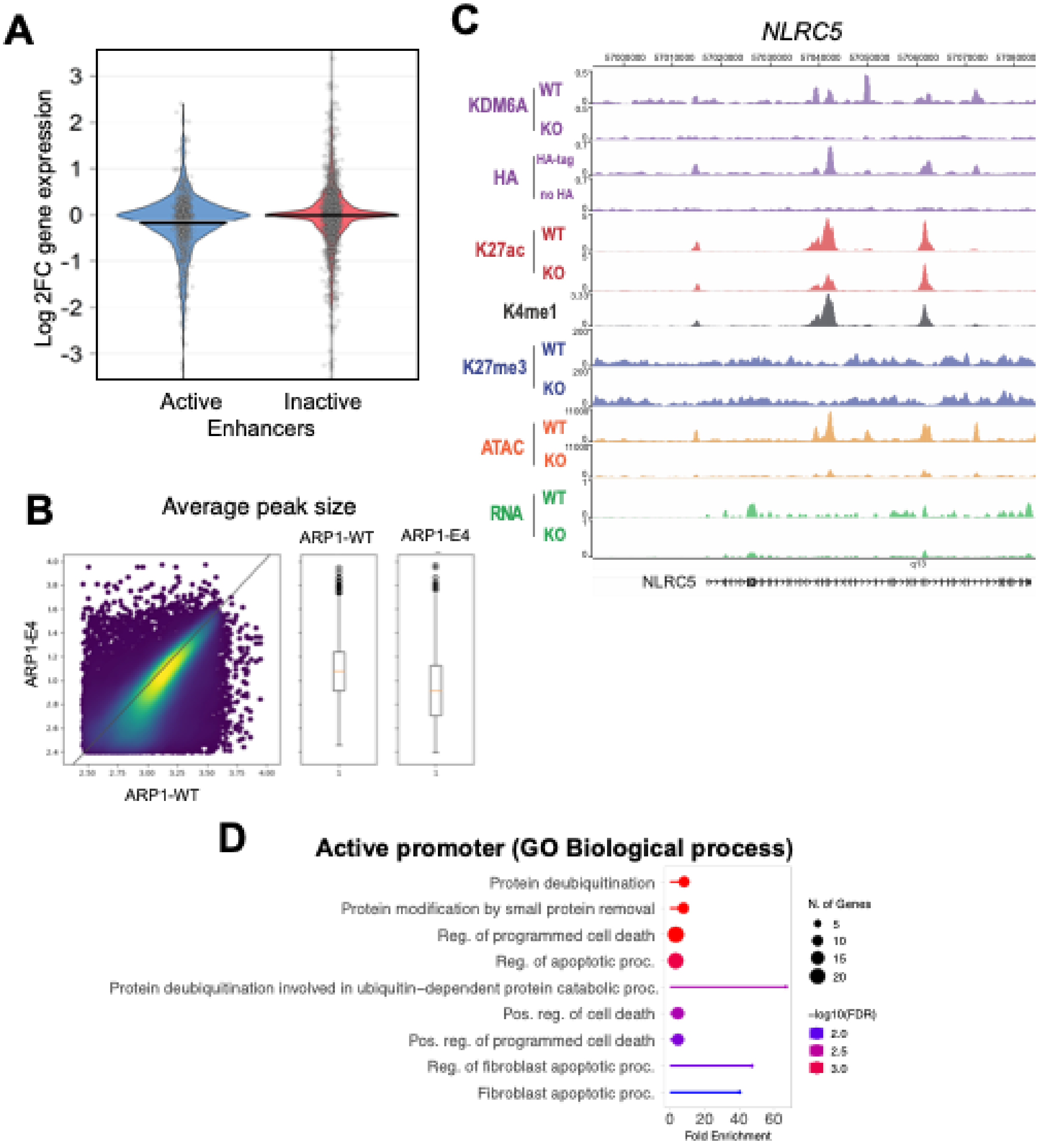


**Figure S6: Epigenetic changes caused by KDM6A loss. A)** Pirate plot show the distribution and mean of gene expression fold change between KO and WT KDM6A at genes associated with either active or inactive enhancers. **B)** ATAC-seq peak size distribution in ARP-1 WT versus E4 KDM6a knockout cells. **C)** Genome browser view of the NLRC5 locus in ARP1 cell lines WT or KO for KDM6A. **D)** ShinyGO pathway enrichment analysis of gene associated with inactive enhancers, promoters, active and inactive or poised enhancers bound by KDM6A.

Figure S7

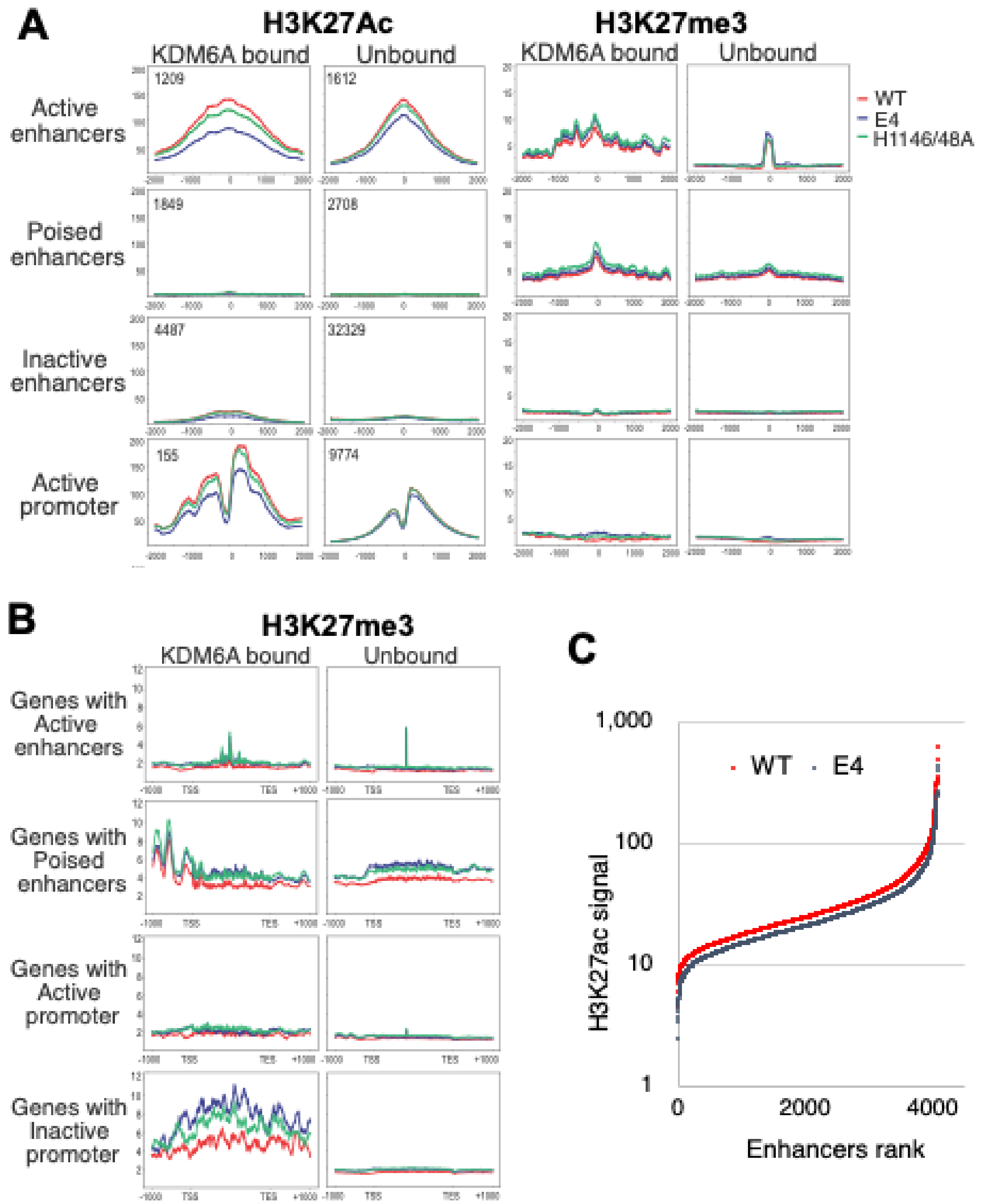


**Figure S7: Epigenetic changes caused by jmjD dead KDM6A. A)** H3K27ac-seq and H3k27me3-seq signals in ARP1 isogenic clonal cell line with KDM6A wild type, KO or mutated at H1146/48 centered on H3K4me1 peaks for enhancers and centered on transcription start sites for promoters **B)** H3K27me3 metagene analysis at locus bound by KDM6A in ARP1 isogenic clonal cell line with KDM6A wild type, KO or mutated at H1146/48. **C)** Hockey stick plots showing H3K27ac signals rank-ordered enhancers in ARP1 WT or KO for KDM6A.

Figure S8

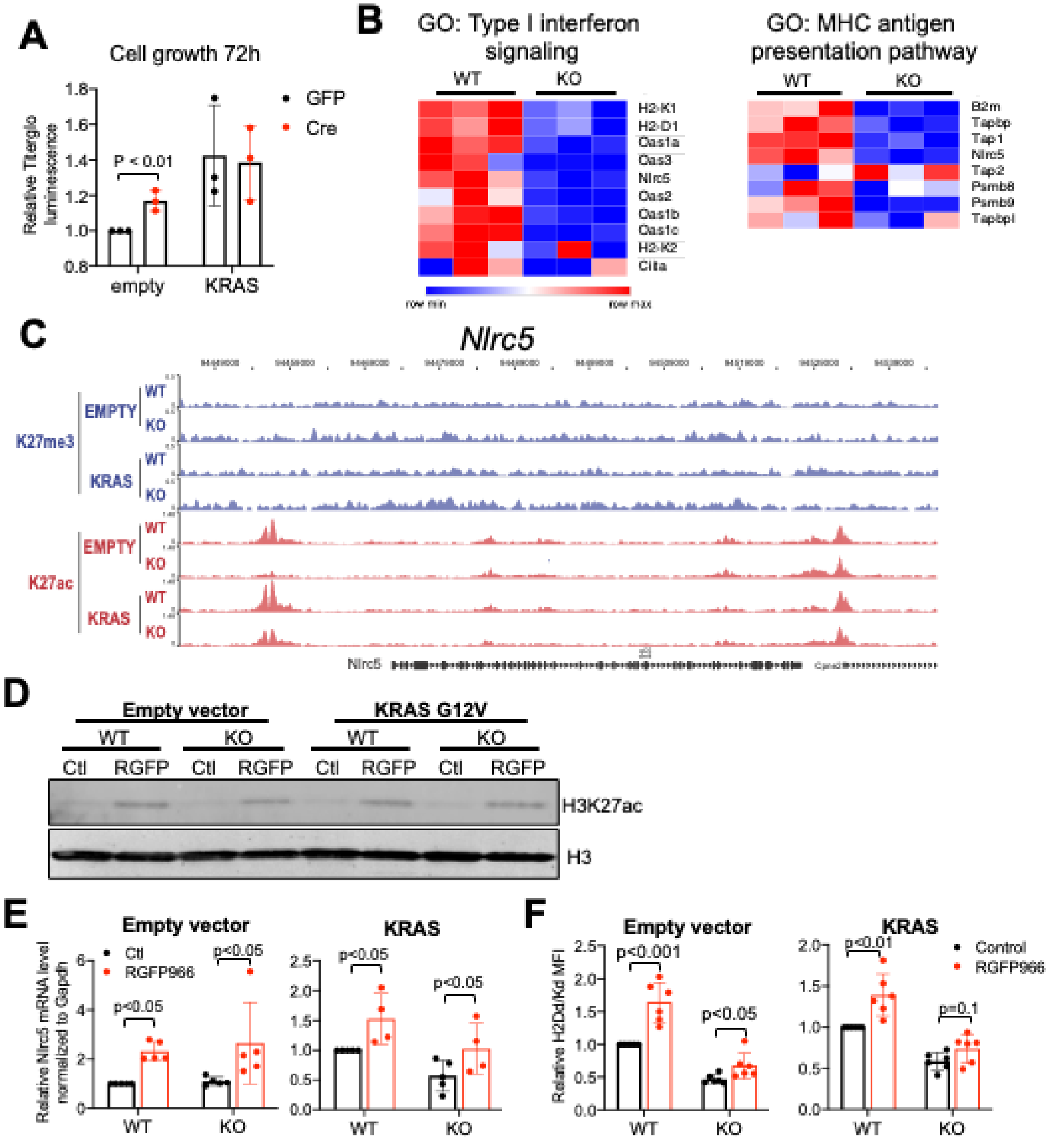


**Figure S8: Phenotypes observed in Kdm6a depleted MEF A,** Relative cell titer-glo fluorescence 72h after plating cells at same density. **B,** Heatmap of expression level (CPM values) of genes included in the indicated GO term in MEF cell lines wild type and KO for *Kdm6a*. **B**, Genome browser view of the CIITA locus in ARP1 cell line WT or KO for KDM6A. **C,** Immunoblot of H3K27ac 72h post treatment with 10µM RGFP966. **D,** mRNA analysis by QPCR normalized to *Gapdh* 72h post treatment with 10µM RGFP966 (4 or 5 biological replicates; +/- SD. Wilcoxon T-test). **E,** MFI quantification of MHCI H2Dd/Kd (5 or 6 biological replicates; +/- SD, Mann-Whitney T-test).

Figure S9

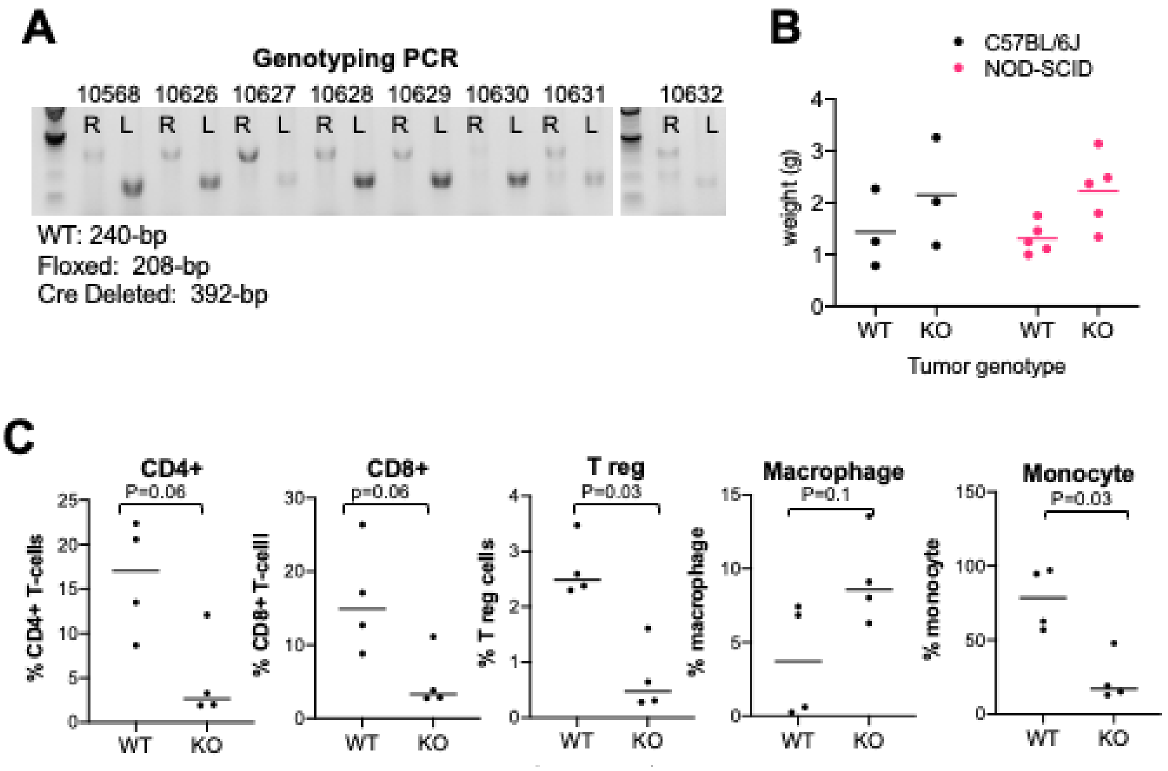


**Figure S9: MEF xenografts experiment A)** Gel electrophoresis of genotyping PCR in xenograft tumors isolated from C57BL/6J mice. **B)** Ras-driven tumors derived from MEFs xenograft tumors were harvested and weighed 3 weeks after injection of tumor cells into C57BL/6J or NOD-SCID mice. **C)** Flow cytometry quantifications of immune cell populations in the xenograft tumors isolated from 57BL/6J in B, 4 biological replicates, Mann Whitney T-test).
